# Tubulin autoregulation mediator TTC5 regulates neuronal morphology and migration

**DOI:** 10.64898/2026.06.23.733857

**Authors:** Stephanie L. Sarbanes, Jakub Ziak, Alex L. Kolodkin, Antonina Roll-Mecak

## Abstract

Microtubule dynamics regulation is critical for neuronal development, yet how neurons regulate tubulin levels remains poorly understood. Tetratricopeptide repeat domain 5 (TTC5) mediates the co-translational degradation of tubulin transcripts in response to excess soluble tubulin, a.k.a. “tubulin autoregulation”, and TTC5 mutations are associated with cerebral atrophy, speech and motor impairment. Despite clinical relevance, the role of TTC5 and tubulin autoregulation in neurons has not been established. Using human induced pluripotent stem cells (iPSCs)-derived cortical-like neurons, we demonstrate that tubulin autoregulation is active in neurons and fully TTC5-dependent. Loss of TTC5 function suppresses microtubule polymerization and impairs axonal outgrowth and arborization, while enhancing cellular motility. These phenotypes recapitulate *in vivo* where TTC5 loss in mouse cerebral cortex projection neurons disrupts axon and dendrite arborization and drives aberrant hypermigration. Patient disease mutants phenocopy this motility dysregulation, directly linking TTC5 dysfunction to clinically relevant neurological deficits. Together, our findings establish tubulin autoregulation and its mediators as essential regulators of neuronal morphogenesis and connectivity and provides a mechanistic framework for understanding TTC5-associated neurodevelopmental disease.

## Introduction

The compartmentalized morphology of neurons as well as their complex stereotyped spatial organization within the brain is controlled by the microtubule cytoskeleton. During neuronal differentiation, transcriptional upregulation of a broad repertoire of α- and β-tubulin isoforms drives rapid assembly of microtubules required for early neurite outgrowth and polarization of a long, extended axon and a dendritic arbor^1,2^. Once established, microtubules serve as both structural scaffolds and tracks for long-range intracellular transport of cargoes pivotal to both the plasticity and the long-term maintenance of neuronal networks. Early neuronal differentiation also encompasses a stage of stereotyped neuronal migration in which newly generated neurons travel from their birthplace to populate their target cortical layer before extending additional processes and forming functional connections^3^. Given these diverse and indispensable roles, microtubule assembly is tightly controlled, including through the availability of the building blocks required for microtubule polymerization, the α/β-tubulin heterodimers.

Classic studies showed that cells buffer changes in their pool of soluble α/β-tubulin heterodimers through a homeostatic feedback response called “tubulin autoregulation” whereby a sudden excess of tubulin, such as upon drug-mediated microtubule depolymerization, deflagellation or change in cell shape, triggers rapid degradation of tubulin transcripts ^4–7^. This turnover occurs co-translationally and is mediated by TTC5 ^8^. When soluble tubulin is in excess, TTC5 marks tubulin-translating ribosomes by recognizing the first four amino acids of β-tubulin (MREI) as they emerge from the ribosome exit channel (or MREC in the case of α-tubulin)^9–11^. TTC5 then binds an adaptor protein, S-phase Cyclin A Associated Protein (SCAPER), which recruits the CCR4-Not deadenylase complex to degrade the tubulin transcript ^8,12–14^ (Figure 1A). Identification of TTC5 and SCAPER provided long-sought molecular handles to dissect the physiological consequences of defects in tubulin autoregulation. Mutations in both *TTC5* and *SCAPER* lead to an intellectual disability syndrome characterized by cerebral atrophy, hypotonia and severe speech and motor impairment suggesting that tubulin autoregulation is particularly critical for nervous system development^15–20^. Notably, these clinical features overlap with those of the many neurodevelopmental disorders tied to mutations in tubulin genes or in genes that alter microtubule stability or structure ^21,22^. Yet, despite the discovery of tubulin autoregulation almost half a century ago, whether this process is operational in neurons, or any other terminally differentiated cell, and how it shapes neuronal development remains unknown.

**Figure 1.**
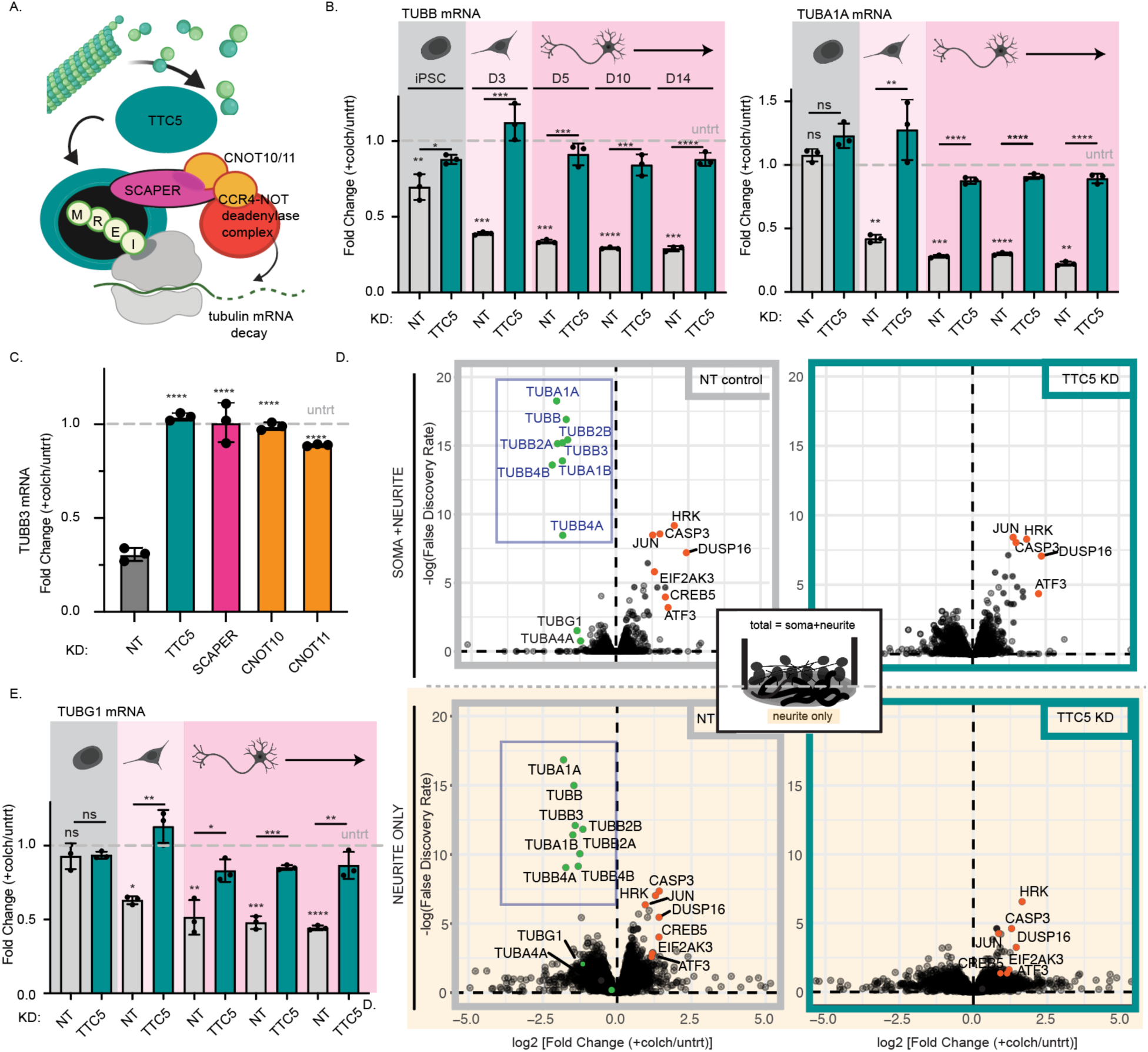
Tubulin autoregulation is active in neurons and relies on TTC5 and other canonical tubulin autoregulation pathway players. (**A**) Graphical depiction of tubulin autoregulation—upon microtubule depolymerization, TTC5 binds to the N-terminal MREI motif of β-tubulin and recruits SCAPER which in turn recruits the RNA decay complex CCR4-Not *via* the CNOT10/CNOT11 module, resulting in co-translational tubulin transcript decay. (B) Fold change for TUBB and TUBA1A mRNA levels upon microtubule depolymerization with colchicine (“+colch”; 10 μM for 5 hours) relative to the untreated (“untrt”) control upon NT (gray) and TTC5 KD (teal) across differentiation from iPSCs to D14 i^3^Neurons assayed by RT-qPCR. Data normalized to housekeeping gene hypoxanthin phosphoribosyltransferase 1 [HPRT]). n = 3 independent replicates, error bars, mean +/-SD; *, p < 0.05, ** p, < 0.01, *** p, < 0.001, **** p, < 0.0001 by multiple unpaired t-test (two-stage step-up method) for both the untreated control and for TTC5 KD relative to NT at each stage. (**C**) Fold change for TUBB3 mRNA levels, quantified as in (B), upon colchicine treatment in D14 i^3^Neurons in TTC5 (teal), SCAPER (magenta), CNOT10 (orange) or CNOT11 (orange) KDs. n = 3 independent replicates, error bars, mean +/-SD ; ****, p < 0.0001 by one-way ANOVA to NT control with Holm-Sidak’s multiple comparisons test. (**D**) Volcano plot of transcriptomic changes upon colchicine treatment of D14 NT (left, gray) and TTC5 KD (right, teal) i^3^Neurons relative to the untreated control in total transcriptome (top) and neuritic transcriptome (bottom) mechanically fractionated via Boyden chamber; tubulin genes labeled by name and marked in green; select stress response genes labeled by name and marked in orange. (**E**) Fold change for TUBG1 mRNA levels upon microtubule depolymerization with colchicine as in (B).

Employing an iPSC-derived human neuron “i^3^Neuron” model, we show that TTC5 loss and TTC5-mediated autoregulation defects impair microtubule dynamics and decrease neurite outgrowth and arborization. By developing a novel glia-guided neuronal migration assay utilizing iPSC-derived neurospheres, we further discover that TTC5 loss drives hypermotility, a phenotype that is recapitulated in non-neuronal lines and upon loss of the TTC5 interactor SCAPER, supporting a broader role for tubulin autoregulation in cellular motility. Complementing these in vitro studies, we present the first evaluation of TTC5 function in a mouse model of cortical development, uncovering context-dependent defects in neuronal arborization and migration. Finally, using a CRISPR knock-in approach, we show that *TTC5* missense mutations identified in patients with intellectual disability syndrome abolish autoregulation and recapitulate the TTC5 knock-down phenotype, directly linking TTC5 dysfunction to disease. Together, this work provides the first characterization of tubulin autoregulation in the polarized context of the neuron, illuminates the cellular and molecular basis of TTC5-associated intellectual disability, and provides a robust panel of i^3^Neuron-based assays capable of capturing clinically relevant neurodevelopmental phenotypes.

## Results

### Tubulin autoregulation is active in neurons but not iPSCs

Tubulin autoregulation is active in a range of cell types, from common immortalized cancer cell lines to primary fibroblasts^6,23^; however, it has yet to be documented in the post-mitotic, polarized neuron. Neurogenesis requires dramatic upregulation of tubulin transcription and translation to drive rapid extension of microtubule-scaffolded neurites ^1,24^. Both TTC5 and SCAPER protein expression track with tubulin upregulation during early differentiation, suggesting that these regulatory processes might be coordinately controlled^25^ (Figure S1A). We therefore examined tubulin autoregulation responsiveness in neurons across this early neurodevelopmental window. We employed a human iPSC-derived neuronal cell culture model engineered with doxycycline(dox)-inducible neurogenin-2 (NGN2) to drive stereotyped differentiation into neurons that have characteristics of glutamatergic excitatory cortical neurons (i^3^Neurons)^26,27^. We then used CRISPR interference (CRISPRi) to knock-down autoregulation mediator TTC5 and confirmed its reduction at both the RNA and protein level at the iPSC stage and upon differentiation into i^3^Neurons (Figure S1B, S1C) relative to a non-targeting (NT) control^28^. Notably, these TTC5 knock-down (KD) lines exhibit no gross differences in iPSC morphology, proliferative ability (Figure S1D) or in gross capacity to differentiate into neuronal progenitor cells (NPCs) and neurons. We acutely triggered tubulin autoregulation by treating cells with the microtubule-depolymerizing drug colchicine^5,6,12^, and then assayed changes in tubulin isoform *TUBB* and *TUBA1A* levels across early differentiation, from iPSCs to day 14 (D14) i^3^Neurons.

Unexpectedly, we find that tubulin autoregulation is not operational at the iPSC stage but active once differentiated into NPCs (D3). While at the iPSC stage, cells treated with colchicine exhibit mild or no reduction in *TUBB* and *TUBA1A* mRNA levels, by D3 post-differentiation colchicine treatment results in ∼ 65% reduction of tubulin mRNAs and is saturated to ∼ 25% of the tubulin mRNAs levels in untreated cells by D5 (Figure 1B). TTC5 knock-down completely abrogates this response (Figure 1B) as does knock-down of downstream tubulin autoregulation pathway components SCAPER and the CCR4-Not complex factors CNOT10 and CNOT11 (Figure 1C), which link TTC5 to the deadenylation complex through SCAPER (Figure 1A, S1E, S1F).

In contrast to the cancer cell lines in which tubulin autoregulation has been so far studied, neurons express a broader repertoire of tubulin isoforms, including neuron-specific TUBB2A, TUBB2B and TUBB3^1^ (Figure S1G). Neurons therefore pose an ideal context for assessing potential isoform-specific differences in autoregulation responsiveness. To better assess tubulin autoregulation across tubulin isoforms, and to obtain a global picture of the impact of TTC5-mediated autoregulation on neuronal transcriptomes, we performed 3’ RNA-Seq on D14 neurons with or without colchicine treatment. This comprehensive analysis revealed the remarkable specificity of the tubulin autoregulatory response. Tubulin isoforms were overwhelmingly the dominant set of downregulated transcripts, with all expressed tubulin isoforms responsive to tubulin autoregulation (Figure 1D [top]). The reduction in β-tubulin (*TUBB*, *TUBB2A*, *TUBB2B*, *TUBB3*, *TUBB4A* and *TUBB4B*) and α-tubulin (*TUBA1A* and *TUBA1B*) transcripts ranges from 3 to 5-fold, with the strongest reduction in *TUBB4B* (∼4.6-fold) and *TUBA1A* (∼4.1-fold). Their coordinated degradation is consistent with recognition by TTC5 of the N-terminal MREC or MREI motifs shared across all α- and β-tubulin isoforms, respectively, ensuring the preservation of isoform ratios which have been shown to possess different polymerization characteristics^29,30^. This response is absent in TTC5 KD cells (Figure 1D [top]). The γ-tubulin *TUBG1* transcript is also reduced (∼2.2-fold) upon colchicine treatment, even though it lacks the MREI motif, which we further corroborated upon colchicine treatment across i^3^Neuron differentiation by qPCR (Figure 1E). Other significantly upregulated transcripts are primarily well-known stress-responsive or pro-apoptotic genes such as *Jun* proto-oncogene [*JUN*]^31–33^, eukaryotic translation initiation factor 2α kinase 3 [*EIF2AK3*, also known as *PERK*]^34^, Harakiri [*HRK*]^35^, Caspase 3 [*CASP3*]^36^, cAMP responsive element binding protein 5 [*CREB5*]^37^ and activating transcription factor 3 [*ATF3*]^38^ ^33^(Figure 1D), a response that is TTC5-independent. Given compartment-specific tubulin transcript localization and translation in neurons^39–41^, we investigated whether tubulin autoregulation extends to neurites as well. By culturing neurons on microporous transwell membranes that restrict cell bodies to the upper chamber while permitting neurite extension into the lower compartment, we could mechanically fractionate neurites from the lower membrane surface to obtain neurite-only (encompassing both proximal and distal neurite) transcriptomes with or without colchicine treatment. This confirmed that tubulin transcripts present in neurites are similarly subject to TTC5-mediated autoregulation (Figure 1D [bottom]). Taken together, these experiments demonstrate that tubulin autoregulation is active in neurons, dependent on TTC5 and other known autoregulation mediators and constitutes the dominant transcriptomic response to manipulation of the soluble tubulin pool in both soma and neurites.

### TTC5 loss impairs arborization and axonal outgrowth

Recent human genetics studies connect TTC5 mutations to intellectual disability. Patients exhibit cerebral atrophy, hypotonia and severe speech and motor impairment from a young age, suggesting a critical role for TTC5 in cortical neurodevelopment^18^. These disease manifestations may reflect disruption of TTC5-dependent processes intrinsic to neurons. Though we observed no gross differences in morphology or capacity to differentiate upon TTC5 KD in the i^3^Neurons during our initial assessments of tubulin autoregulation, we reasoned that TTC5 loss might instead impact early morphological features of neuronal differentiation, specifically the length of axonal processes and subsequent process elaboration. To assess morphology, we subcloned from our parental CRISPRi iPSCs a reporter line expressing both diffuse mScarlet as a fluorescent cell fill and EGFP-tagged end binding protein 1 (EB1), which binds to growing microtubule ends and reports on microtubule dynamics ^42^. Using this reporter line, we examined the impact of TTC5 loss on elaboration of the smaller neurites proximal to the soma that constitute the early formation of dendritic arbors. i^3^Neurons seeded onto poly-L-ornithine (PLO)/laminin at low density are slower to arborize and their somas tend to form clusters, hindering assessment of neurite morphology. To visualize individual cell arbors, we seeded the mScarlet-labeled cells on a glial feeder layer that fosters viability and maturation of i^3^Neurons, particularly at low densities ^43^ (Methods). At D12, we quantified arborization parameters, including primary neurite number, neurite complexity and total neurite length (Figure 2A, Figure S2A, Movie S1). Using an automated analysis pipeline to skeletonize branches emanating from a binarized cell body mask (Figure S2B, Movie S1, Methods), we found that TTC5 KD neurons exhibited a significant ∼ 40% reduction in total neurite length per neuron (142+/-77 μm) relative to the NT controls (231 +/- 109 μm) (Figure 2B), driven by reductions both in branch number (complexity) and individual neurite length. To assess neurite complexity - encompassing primary, secondary and the less-common tertiary neurites - we quantified the total number of branch ends *per* neuron. This revealed a significant decrease from an average of 8.7+/-3 total neurites in NT control neurons to 6.3 +/-3 in TTC5 KD neurons (Figure 2B). As a proxy estimate for the number of primary neurites (extending directly from the soma), we subtracted the number of branchpoints from branch ends, which also revealed a consistent reduction in primary neurite number upon TTC5 KD (Figure 2B).

**Figure 2.**
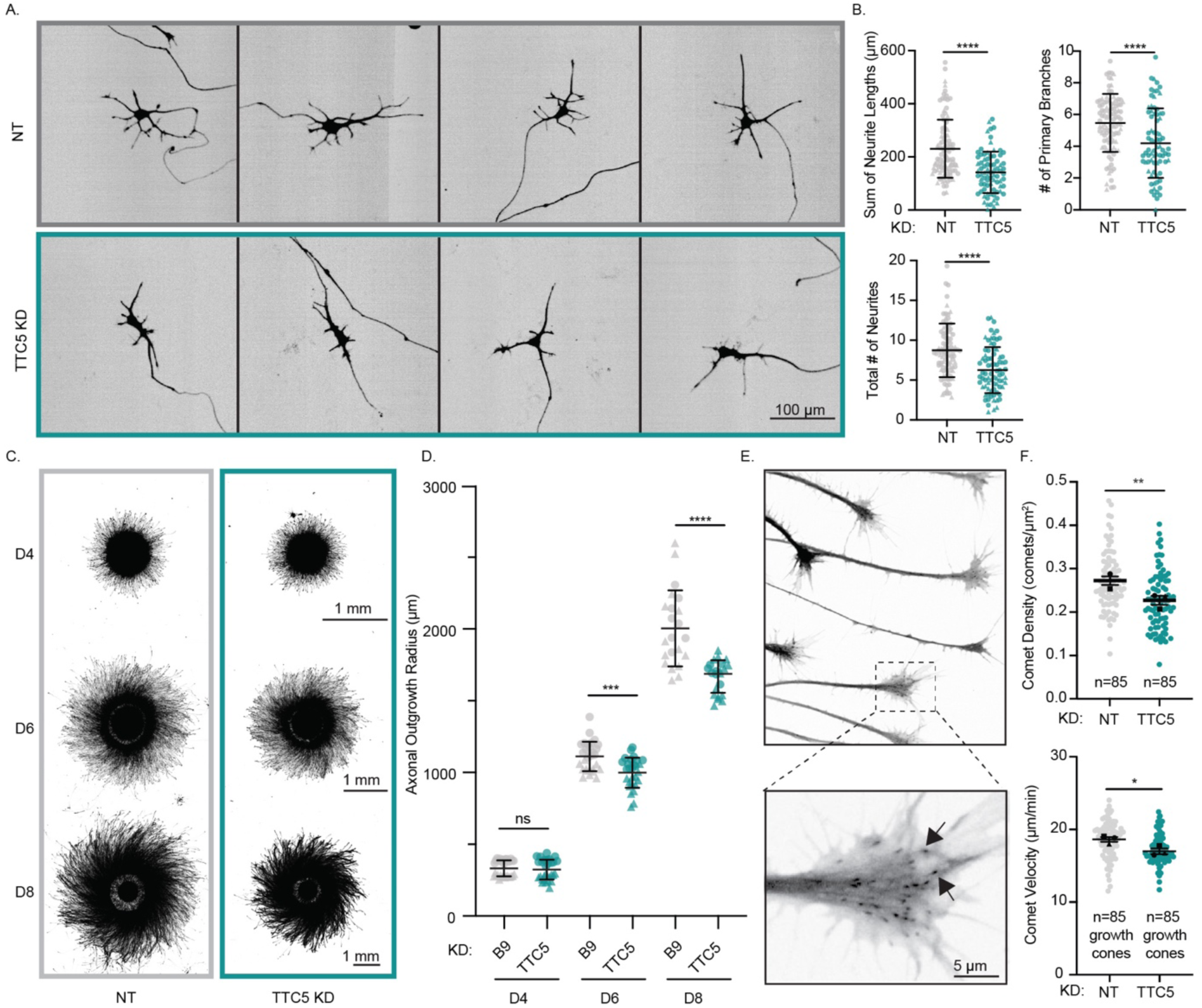
TTC5 loss impairs arborization, axonal outgrowth and microtubule dynamics. (**A**) Representative images of mScarlet-expressing D12 NT or TTC5 KD i^3^Neurons (also see Movie S1); scale bar: 100 μm. (**B**) Arborization metrics: sum of neurite lengths, number of primary branches, and total number of neurites (primary, secondary and tertiary) averaged across 72 timepoints per neuron (Methods). Individual dots represent individual neurons. n = 102, 79 for NT (gray) and TTC5 KD (teal), respectively, from two independent experiments (circle/triangle symbol); error bars, mean +/- SD; ****, p < 0.0001, *, p < 0.05 by Welch’s t-test. (**C**) Neurospheres of mScarlet- and EB1-EGFP-expressing NT and TTC5 KD neurons (Methods) at D4, D6 and D8 post-differentiation; scale bar: 1 mm. (**D**) Axonal outgrowth at each timepoint, quantified as the radius of the thresholded signal normalized to the radius of the inner sphere of cell bodies (Methods). Individual dots represents individual neurospheres; n = 31, 32 (D4), 28, 29 (D6), 22, 22 (D8) neurospheres for NT (gray) and TTC5 KD (teal), respectively, across two independent differentiations (circle/triangle symbol); error bars indicate mean +/- SD; ***, p<0.001, ****, p <0.0001 by Welch’s t-test. (**E**) Representative frame from a time lapse of EB1-EGFP comets in growth cones at the outer edge of a neurosphere (top) with magnification of a single growth cone (bottom); EB1-GFP comets indicated by arrowheads; scale bar: 5 μm. (**F**) Comet density (top) and velocity (bottom) in growth cones of D5 NT (grey) and TTC5 KD (teal) i^3^Neurons (Methods, also see Movie S2). Individual dots represent means from individual growth cones from three independent experiments; n = 85 growth cones per condition; error bars and symbol indicate paired means +/- S.E.M. of each independent experiment, *, p < 0.05, **, p < 0.01 by paired t-test.

We next characterized the impact of TTC5 KD on axon growth. To do so, we differentiated the iPSCs into “i^3^Neurospheres”, spherical clusters of cell bodies and proximal processes of i^3^Neurons which extend neurites in a stereotyped radial arrangement^44^. This geometry enables large-scale assessment of axonal outgrowth from thousands of neurons, quantified as the radius of outgrowth from the sphere boundary (Figure 2C). Prior characterization using compartment-specific staining and EM-based assignment of microtubule polarity of neurons differentiated in the same manner confirmed classification of these longer neurites are axonal^44,45^. TTC5 knock-down significantly reduces axon outgrowth, an effect which is increasingly visible over the course of differentiation (Figure 2C, 2D). Outgrowth radius was comparable between conditions at D4 (333.00 +/- 56 μm for NT versus 324 +/- 69 μm for TTC5 KD), but by D6, TTC5 KD neurons showed a 10% decrease in outgrowth radius, which progressed to a 17% reduction by D8 (2004 +/- 265 μm for NT versus 1669 +/-113 μm for TTC5 KD) (Figure 2D). Together these findings establish that TTC5 loss impairs the establishment of polarized neuronal morphology, reducing both axon extension and dendritic arborization.

### TTC5 loss impairs microtubule growth

The rapid extension of neuronal processes relies upon a dynamic microtubule cytoskeleton with each growing process driven by microtubule polymerization. Consistent with this, manipulation of microtubule dynamics directly impacts neurite outgrowth^46–49^. Using our EB1-EGFP reporter, we tracked microtubule plus-end comets within the growth cones at the distal tips of neurites at the edge of the i^3^Neurospheres (Methods^50^) (Figure 2E, Movie S2). TTC5 KD neurons showed a ∼17% reduction in EB1 comet density relative to the control, indicating reduced microtubule polymerization (Figure 2F), accompanied by a modest ∼9% reduction in microtubule growth velocity (Figure 2F, see Figure S3A for distribution of individual comet velocities within each growth cone). Notably, while tubulin transcript levels are slightly reduced in TTC5 KD neurons (Figure S1G, S3B), there is no large change in global tubulin protein levels within the precision limit of Western blot-based detection (Figure S3C, S3D). To determine whether this effect on microtubule dynamics is more general, we imaged microtubule dynamics in EB1-mNG knock-in U2OS cells upon acute siRNA knock-down of TTC5 (Figure S4A, S4C). TTC5 KD U2OS cells display significant reductions in comet density (∼30%) and comet velocity (∼15%) (Figure S4B), consistent with a recent report examining TTC5 knock-out in HeLa cells^51^ and establishing reduced microtubule growth as a generalizable feature of TTC5 loss. Taken together, these data indicate that TTC5 loss impairs arborization and axonal outgrowth in concert with a reduction in microtubule growth dynamics. These observed defects in microtubule dynamics run counter to the more naive prediction for the impact of TTC5 knock-down: by removing the negative feedback of tubulin autoregulation, TTC5 knock-down might have been expected to increase tubulin transcript levels, elevate soluble tubulin levels, and thereby enhance microtubule growth and neurite extension. That the opposite occurs, suggests a more complex relationship between autoregulatory feedback, homeostatic tubulin levels, and microtubule dynamics.

### TTC5 regulates neuronal motility

Following early neuronal differentiation and initial neurite outgrowth, cortical neurons born in the subventricular zone (SVZ) migrate radially outward along a scaffold of radial glia to populate their appropriate neocortical laminar position in the cortical plate (CP) ^3^. Spatiotemporal control of this process is critical for subsequent establishment of functional connectivity through axon and dendrite elaboration. Disruption in this migratory process leads to clinical manifestations such as lissencephaly ^3^ and has in some cases been correlated with altered neurite branching ^52,53^. To assess a role for TTC5 in neuronal migration, we developed an in vitro migration assay by replating the neurospheres on a monolayer of glia instead of the traditional PLO/laminin substrate. In contrast to neurospheres cultured on PLO/laminin, adherence to the glia monolayer enhances exit of neuronal cell bodies from the central sphere and into the imaging plane, enabling tracking of radial nuclear trajectories using a fluorescent Halo-NLS (nuclear localization signal) reporter. Using a bulk dispersion assay in which high-percentage Halo-NLS-labeled neurospheres were monitored over an early differentiation timecourse, we observed that TTC5 loss leads to a dramatic increase in nuclear dispersion relative to the NT control (Figure 3A). To quantify this, we generated radial profiles of nuclear signal intensity from the sphere center outward, binning signal into “inner”, “edge” and “outer” zones (Figure 3B, right panel). While NT and TTC5 KD distributions were similar at D6, by D9 and D12, TTC5 KD neurospheres showed substantially elevated signal in both the inner and outer zones (Figure 3B, S5A), accompanied by a commensurate decrease in signal at the central sphere edge and consistent with enhanced displacement of the TTC5 KD nuclei away from their starting positions (Figure 3B, S5A). Loss of TTC5 therefore drives enhanced neuronal dispersion.

**Figure 3.**
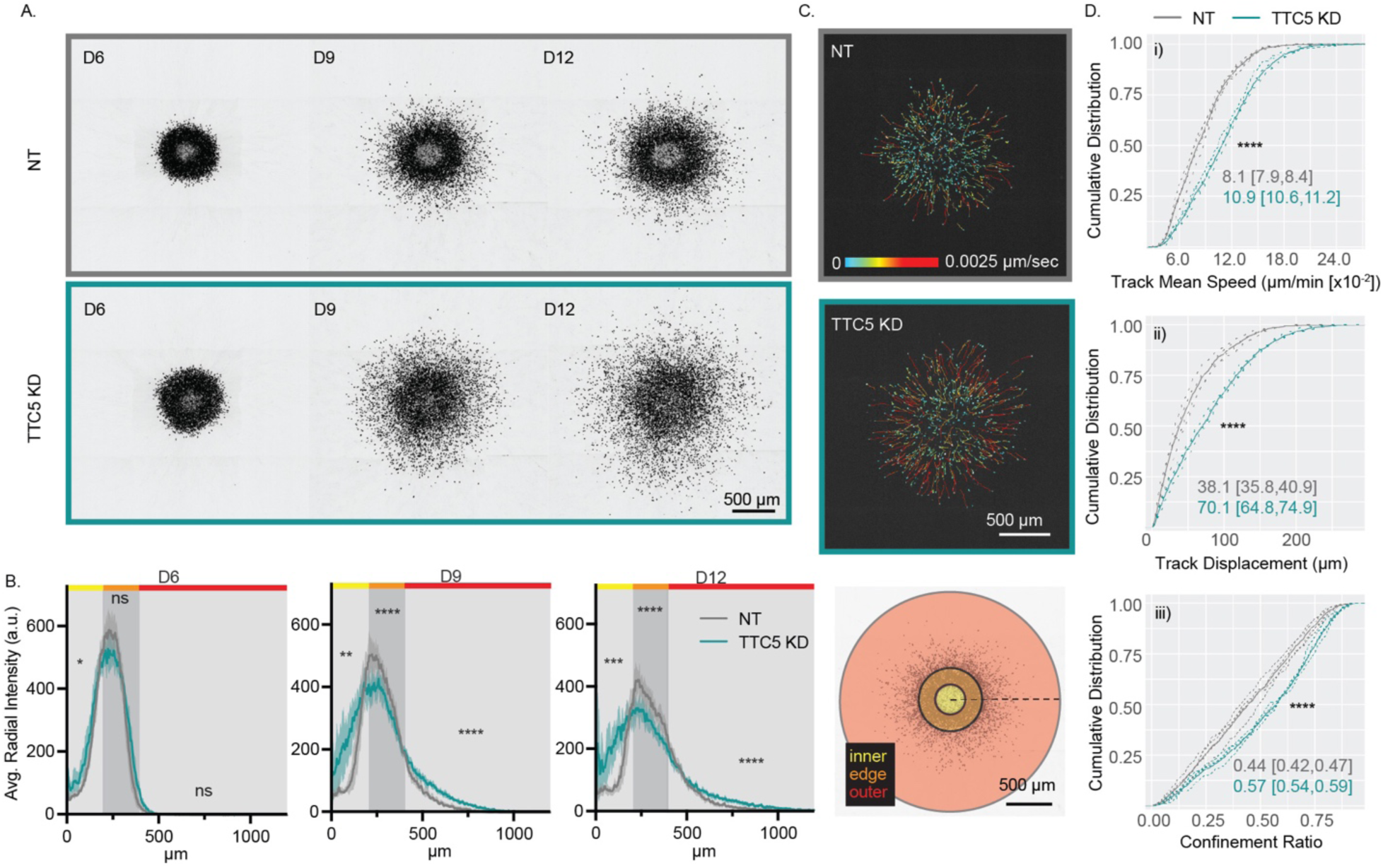
TTC5 loss results in hypermotility in a glia-guided neuronal migration assay. (**A**) Representative images of Halo-NLS-expressing NT control and TTC5 KD neurospheres at D6, D9 and D12 post-differentiation showing nuclear distribution over time (maximum intensity projections). Scale bar: 500 μm. (**B**) Nuclear migration profiles at each timepoint quantified as the averaged intensity of a line extending from the sphere center to the periphery (black dotted line in graphic in rightmost panel) and rotated radially. Each profile depicts the mean (solid line) and S.D. (shadow) across n = 8 neurospheres for both the NT (gray) and TTC5 KD (teal) condition, respectively. Statistical analysis performed using the binned signal intensity from the designated “inner” (yellow), “edge” (orange) and “outer” (red) regions of sphere for each timepoint (see Figure S5A for additional details); *, p < 0.05, **, p < 0.01, ***, p < 0.001, ****, p < 0.0001 by unpaired t-test. (**C**) Representative images of low-percentage NT or TTC5 KD Halo-NLS-expressing neurospheres overlaid with track trajectories color-coded by track mean speed (also see Movie S3). Scale bar: 500 μm. Tracks were analyzed for motility parameters shown in (D). (**D**) Cumulative distribution functions (CDF) for i) track mean speed, ii) track displacement and iii) confinement ratio (a metric for efficiency of motion as the ratio of displacement over total distance traveled) with dotted lines representing profiles for individual neurospheres and solid lines their average; n = 4 NT, 4 TTC5 KD neurospheres comprising 1653 and 1397 tracked nuclei respectively; median and 95% confidence interval indicated on graph and calculated based on corresponding scatterplots in Figure S5B; ****, p < 0.0001 by Kolmogorov-Smirnov test.

To characterize this motility phenotype at single cell resolution, we generated neurospheres with sparse Halo-NLS expressing cells (7.5%) and tracked individual nuclei every 15 minutes for 24 hours from D8 to D9 post-differentiation (Figure 3C, Movie S3, Methods). TTC5 KD neurons displayed a ∼ 33% increase in median track mean speeds (the average of the instantaneous velocities of all points in a track) and enhanced directedness of motion (the ratio of net displacement over total distance traveled), yielding a ∼1.8-fold increase in median net displacement (Figure 3D, Figure S5B). Because TTC5 KD neurons comprised a small fraction of the overall population of neurons within these neurospheres, this increased motility is unlikely to reflect emergent properties of collective migration or differences in overall neurosphere cell packing and adhesion. Rather, it points to an intrinsic cell-autonomous difference in the motile capacity of TTC5 KD neurons. Furthermore, tracking of sparsely-plated individual Halo-NLS-labeled HEK293 cells on a fibronectin substrate also displayed increased motility parameters upon TTC5 loss (Figure S6A, S6B), demonstrating that TTC5-dependent regulation of cell motility is not neuron or substrate-specific. Taken together, these findings reveal striking increases in neuronal dispersion and motility upon TTC5 loss and identify TTC5 as a regulator of cell motility across cell types.

### Autoregulation-defective TTC5 mutant phenocopies morphology and migration defects

Since tubulin autoregulation is active in neurons (Figure 1B), we asked whether the neuronal phenotypes we observe upon TTC5 KD reflect the specific loss of TTC5 function in tubulin autoregulation. To test this, we leveraged a structure-guided mutation, R147A ^8^. R147 forms a salt bridge with the glutamate in the MREI repeat at the N-terminus of β-tubulin (Figure 4A); mutation of the arginine to alanine abolishes binding to the tubulin N-terminus and renders TTC5 unable to mediate tubulin autoregulation (Figure 4A). Using CRISPR knock-in (KI), we analyzed two clonal lines (R147A KI #1 and #2) with the R147A mutation introduced into the endogenous *TTC5* locus (Figure S7A). R147A KI mutation completely abolished tubulin autoregulation over a timecourse of early differentiation (Figure 4B, Figure S7B). The R147A KI mutants phenocopied TTC5 KD across all morphological and microtubule dynamics readouts: axonal outgrowth was reduced in our neurosphere assay (Figure 4C, 4D), EB1 comets density and microtubule growth rates in axonal growth cones were reduced (Figure 4E, 4F, Figure S8A), and arborization metrics of summed neurite length and total neurite number were also reduced (Figure 4G). The R147A KI mutant neurons also recapitulated the hypermotility phenotype observed upon TTC5 KD in both bulk dispersion and single-cell tracking assays (Figures 4H, 4I and S8B, Movie S4). As upon TTC5 KD, baseline tubulin transcript levels were lower in the R147A KI neurons compared to WT (∼50%; Figure S7C), while global tubulin protein levels were similar between the WT and the KI neurons (Figure S7D).

**Figure 4.**
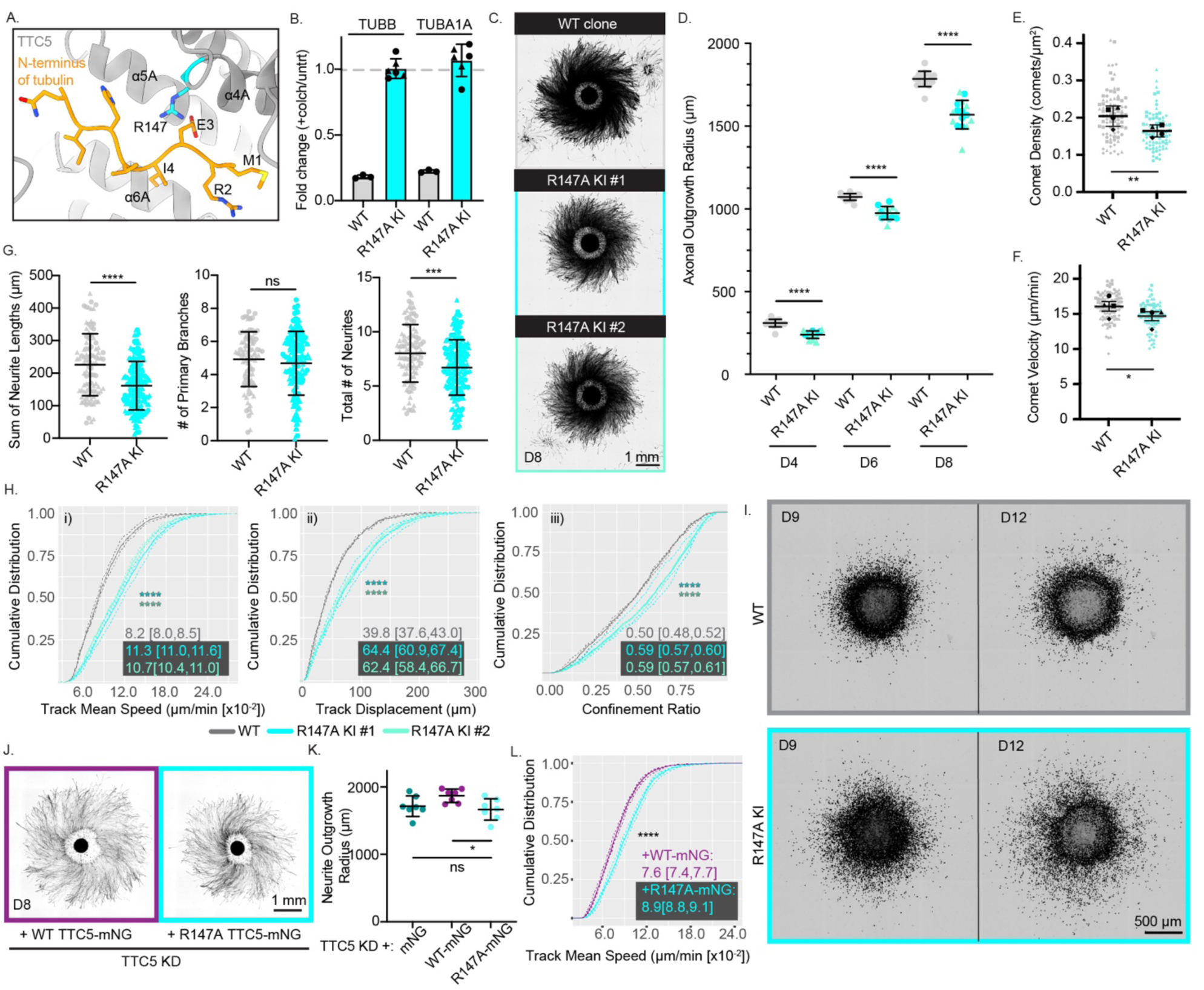
Autoregulation-defective TTC5 mutant R147A phenocopies TTC5 KD morphology and motility defects. (**A**) Structure of the TTC5 pocket (PDB 6T59^8^), highlighting R147 (cyan) whose mutation to alanine abolishes binding of the MREI tubulin motif ^8^. (**B**) RT-qPCR for TUBB and TUBA1A mRNA expression in WT (grey) and R147A KI clone # 1 (cyan) upon colchicine treatment (Methods) in D14 i^3^Neurons (fold change relative to the untreated control following normalization to HPRT). (**C**) Representative images of mScarlet and EB1-EGFP-expressing WT and R147A KI clone #1 and 2 neurospheres at D8 (maximum-intensity projection). Scale bar: 1 mm. (**D**) Axonal outgrowth for the WT clone and two R147A KI clones at D4, D6 and D8; n = 16 spheres for WT and 8 spheres each for R147A KI clone #1 (circle) and #2 (triangle). Error bars, mean +/- SD; ****, p < 0.0001 by Welch’s t-test. (**E-F**) Average EB1-EGFP comet density (E) and comet velocity (F) per growth cone for WT and R147A KI clone #1 (data for additional clones shown in Figure S8A). Dots represent average values from individual growth cones from four independent experiments; n = 87 and 76 growth cones for WT and R147A KI, respectively; error bars and symbol indicate the paired means of each experiment +/- SEM; *, p < 0.05, **, p < 0.01 by paired t-test. (**G**) Arborization metrics averaged per neuron as in Figures 2A, 2B. Dots represent individual neurons from D12 WT (gray, n = 93), and R147A KI (cyan, n = 188 pooled between R147 KI clone #1 and clone #2) i^3^Neurons from two independent experiments (circle/triangle symbol); error bars indicate mean +/- SD; *, p < 0.05, **, p < 0.01, ***, p < 0.001, ****, p < 0.0001 by Welch’s t-test. (**H**) CDFs for i) track mean speeds, ii) track displacement and iii) confinement ratio of tracked nuclei in glial migration assay for Halo-NLS WT clone (n = 3 neurospheres, 1403 nuclei) and R147A KI clone #1 (n = 2 neurospheres, 1820 nuclei) and R147A KI clone #2 (n = 2 neurospheres, 1264 nuclei) (also see Movie S4); dotted lines represent individual neurospheres, solid lines, combined average; median and 95% confidence interval indicated on graph and calculated based on corresponding scatterplots in Figure S8B with statistical significance calculated via Kruskal-Wallis test; **** p < 0.0001. (**I**) Representative images of Halo-NLS- expressing WT control and R147A KI neurospheres at D9 and D12 showing nuclear distribution over time (maximum-intensity projection). Scale bar: 500 μm. (**J**) Representative images of TTC5 KD neurospheres at D8 expressing WT TTC5-mNG (purple) or R147A TTC5-mNG (cyan) (maximum-intensity projection). Scale bar: 1 mm. (**K**) Axonal outgrowth for neurospheres in (J) with comparative inclusion of TTC5 KD neurospheres expressing mNG alone (teal); n = 7 spheres/condition. Error bars, mean +/-SD; *, p < 0.05 by one-way ANOVA with Tukey’s correction for multiple comparisons. (**L**) CDF for mean speeds of tracked nuclei in glial migration assay for Halo-NLS-expressing TTC5 KD neurons upon reintroduction of either WT TTC5-mNG (purple, n = 4 neurospheres, 3013 nuclei) or R147A TTC5-mNG (cyan, n = 4 neurospheres, 3258 nuclei); dotted lines represent individual neurospheres, solid lines, combined average; median and 95% confidence interval indicated on graph and calculated based on corresponding scatterplots in Figure S8C with statistical significance calculated via Kruskal-Wallis test; **** p < 0.0001.

To independently confirm that the neuronal phenotypes depend specifically on TTC5’s tubulin autoregulation function, we performed rescue experiments in TTC5 KD neurons by reintroducing either WT or R147A mutant TTC5 tagged with mNeonGreen (mNG) after fluorescence-activated cell sorting (FACS) on the mNG signal to ensure comparable expression levels. Reintroduction of WT TTC5 counteracted both the defect in axonal outgrowth (Figure 4J, 4K) and the heightened motility (Figure 4L, S8C), while the R147A mutant failed to do so. Alongside our R147A knock-in phenotypes, these findings indicate that the tubulin autoregulation function of TTC5 is required to maintain normal neuronal morphology and migration.

### Loss of SCAPER recapitulates TTC5 KD morphology and migration defects

To further assess the specific contribution of tubulin autoregulation to the observed neuronal phenotypes, we examined a second autoregulation pathway component, SCAPER ^13^. SCAPER acts downstream of TTC5 recruitment to ribosomes by binding to both TTC5 and the CNOT11 subunit of the CCR4-NOT complex to connect tubulin-translating ribosomes to RNA degradation machinery (Figure 1A) ^13^. Like TTC5, SCAPER expression increases during early neuronal differentiation in our i^3^Neuron model ^19,25^ (Figure S1A). SCAPER knock-down abolishes tubulin autoregulation in i^3^Neurons (Figure 1C) and SCAPER mutations are associated with intellectual disability ^20,54^. SCAPER KD neurons phenocopied TTC5 KD defects in arborization (Figure 5A) and reduced axonal outgrowth (Figure 5B, 5C, Figure S9A). Furthermore, loss of SCAPER recapitulates the hypermotility phenotype observed for the TTC5 KD or the R147A mutant knock-in (Figure 5D, S9B). The convergent phenotypes produced by TTC5 KD, targeted elimination of tubulin autoregulation with the R147A mutation and SCAPER KD together strongly implicate tubulin autoregulation in the morphology and migration defects we observe.

**Figure 5.**
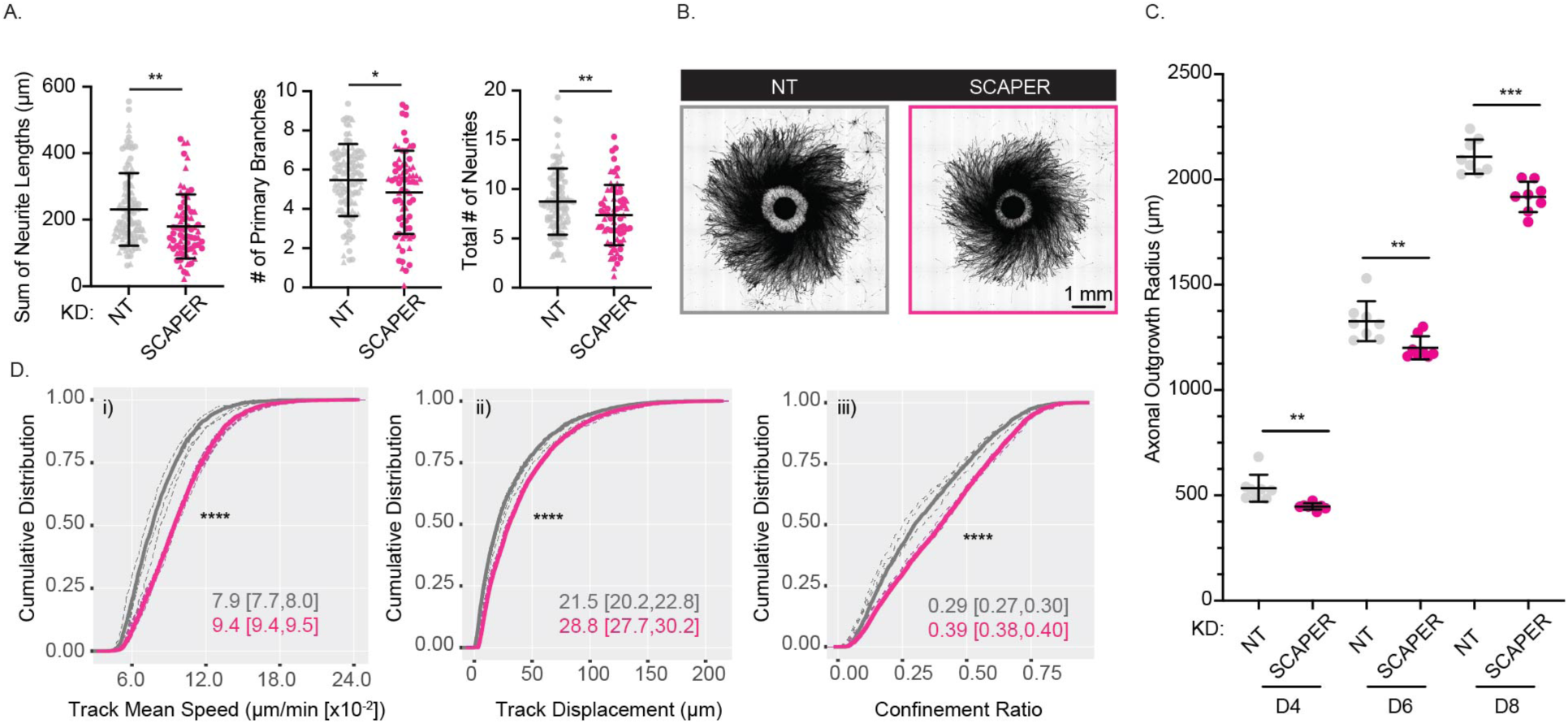
Loss of tubulin autoregulation mediator SCAPER recapitulates TTC5-dependent neuronal phenotypes. (**A**) Quantification of arborization metrics for NT (gray) and SCAPER KD (magenta) neurons (n = 102, 70 neurons for NT and SCAPER KD, respectively) compiled across two independent experiments (circle/triangle symbol). (**B**) Representative images of mScarlet-expressing NT and SCAPER KD neurospheres at D8 (maximum-intensity projected). Scale bar: 1 mm. (**C**) NT (gray) and SCAPER KD (magenta) neurospheres were assayed longitudinally at D4, D6 and D8 post-differentiation for axonal outgrowth (n = 8 spheres NT and 8 spheres SCAPER KD). Error bars indicate the mean +/- SD; ** p<0.01, *** p<0.001 by Welch’s t-test. (**D**) CDF for i) mean speeds, ii) track displacement and iii) confinement ratio of tracked nuclei in glial migration assay for Halo-NLS NT (n = 4 neurospheres, 2249 nuclei) and SCAPER KD (n = 4 neurospheres, 4277 nuclei); dotted line represents individual neurospheres, solid lines, combined average; ****, p < 0.0001 by Kruskal-Wallis test based on corresponding scatterplots in Figure S9B.

### TTC5 disease mutants display aberrant motility

Patients with *TTC5* mutations display a spectrum of clinical features characteristic of an autosomal recessive intellectual disability (ARID) syndrome^18^, yet the impact of specific patient mutations on TTC5 activity remains unknown. Most reported patient mutations result in premature translation termination at various points along the protein, which we predict would be functionally similar to loss of TTC5. However, the effect of missense mutations is less clear. We focused on two missense mutations, A231V and G186V, associated with moderate to severe intellectual disability (Figure 6A, S10A) ^15,16,18^. In patients, the G186V mutation was identified heterozygous to a second allele with a premature stop codon, and individuals harboring this genetic makeup also present with seizures. Mapping these mutations on the structure of TTC5^8^, we predicted that a substitution of A231 to a valine might introduce steric clashes with the backbone of A243 and the sidechain of F247 in the adjacent TPR helix (Figure 6A), while mutation of G186 to a bulkier valine would clash with A209 and Y210 (Figure 6A), in both cases destabilizing the protein. However, given that proteins can accommodate structural perturbations through internal dynamics, especially subtle ones like those predicted for TTC5, functional predictions from *in silico* modeling alone are insufficient. We therefore generated homozygous CRISPR knock-in lines for each mutation, and a heterozygous A231V KI (Figure S10B). Although *TTC5* RNA levels were unchanged (Figure S10C), TTC5 protein signal was undetectable in both homozygous lines (Figure 6B), confirming that these mutations destabilize the protein and trigger its degradation. These findings indicate that patient phenotypes are primarily driven by TTC5 protein loss rather than a toxic gain of function.

**Figure 6.**
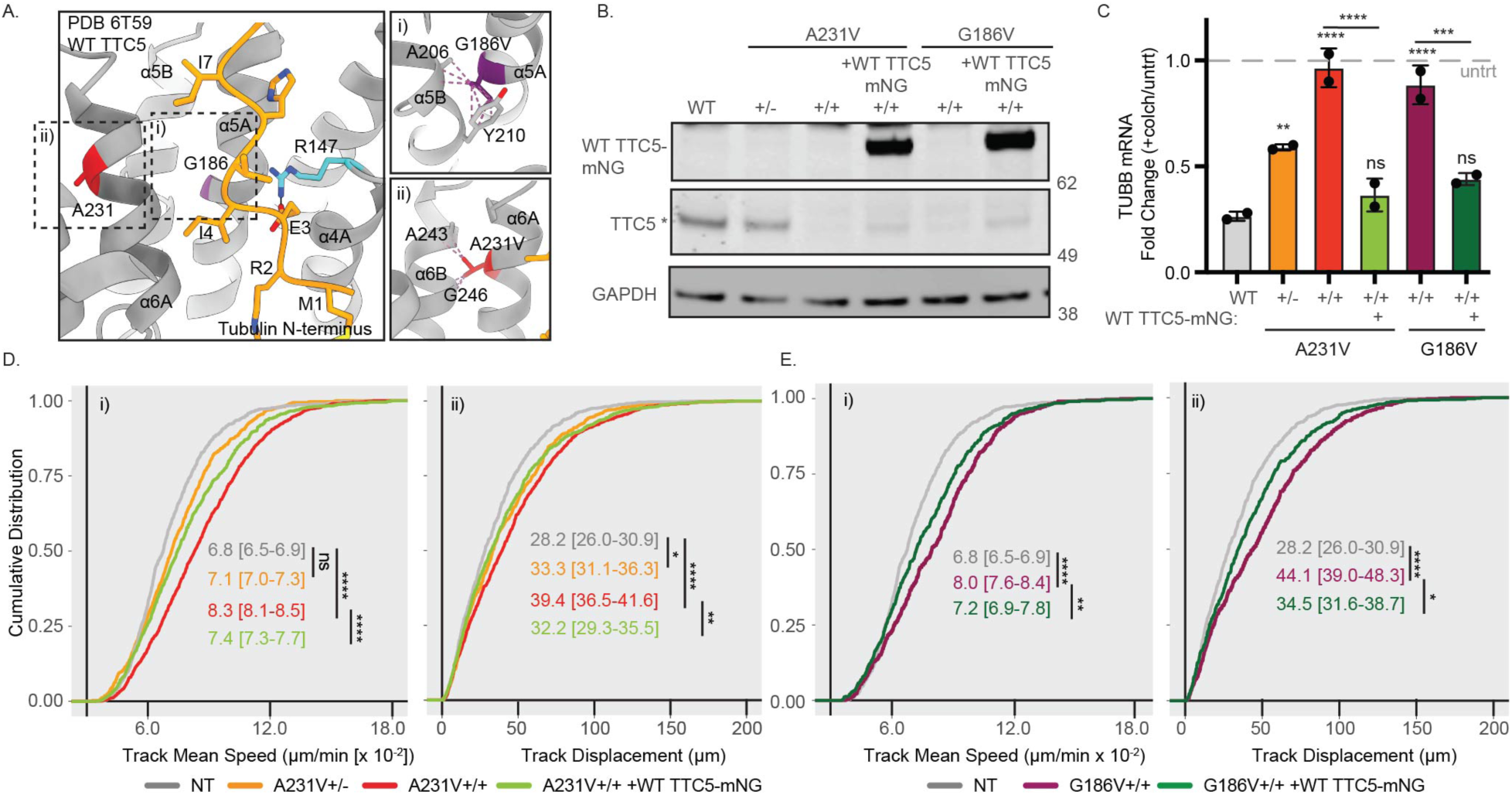
CRISPR knock-in of TTC5 disease mutations results in tubulin autoregulation loss and hypermotility. (**A**) Ribbons structure of TTC5 (PDB ID: 6T59^8^) highlighting location of residues mutated in patients with intellectual disability syndrome: G186 in purple and A231 in red (ball-and-stick representation) in helices α5A and α6A, respectively. The tubulin N-terminal MREI motif is shown in gold and the R147 residue critical for MREI motif recognition shown in cyan (ball-and-stick). (**B**) TTC5 protein expression of both endogenous (*) and WT TTC5-mNG rescue construct, in D10 WT and indicated disease mutant knock-in lines assayed by Western blot (with GAPDH control). (**C**) RT-qPCR for TUBB mRNA expression in WT, disease mutant knock-in and WT TTC5-mNG rescue D10 i^3^Neurons upon acute colchicine treatment (fold change calculated relative to untreated control following normalization to HPRT). n = 2 independent replicates, error bars, mean +/- SD; **, p < 0.01, ***, p < 0.001, ****, p < 0.0001 by ordinary one-way ANOVA with Tukey’s correction for multiple comparisons for each disease mutant and rescue line relative to WT and to each other. (**D-E**) Cumulative density functions for i) mean speed and ii) displacement of tracked Halo-NLS nuclei from (D) WT and indicated A231V disease mutant and WT TTC5-mNG-rescue neurospheres (n = 3 neurospheres each; nuclei = 698 WT, 663 A231V -/+, 1217 A231V +/+, 819 A231V +/+ with WT TTC5-mNG) and (E) G186V disease mutant and WT TTC5-mNG-rescue neurospheres (n = 3 neurospheres each, nuclei = 605 G186V +/+, 523 G186V +/+ with WT TTC5-mNG) with median and 95% confidence interval shown on the graph; *, p < 0.05, **, p < 0.01, ****, p< 0.0001 by Kruskal-Wallis test based on corresponding scatterplots in Figure S10F.

Consistent with complete loss of TTC5 protein, the A231V +/+ and G186V +/+ neurons are completely defective for tubulin autoregulation following acute drug-induced microtubule depolymerization (Figure 6C). Notably, the A231V +/- heterozygous line, expressing half the level of the WT protein, displays a proportionally reduced autoregulation response at ∼50-60% of the WT. In the migration tracking assay, both homozygous mutants display hypermotility compared to the WT control clone, with increased track mean speed and track displacement (Figure 6D, Figure S10F), consistent with what we had observed for the TTC5 KD. The A231V +/- line again displays an intermediate phenotype, with track displacement falling between WT and the homozygous mutant. To rule out clone-to-clone variability in the observed hypermigration, we reintroduced WT TTC5 tagged C-terminally with mNeonGreen (TTC5-mNG) into each homozygous mutant line (Figure 6B). This partially rescued motility in both mutant lines and restored the autoregulation response (Figure 6C, 6D, S10F). As observed for R147A KI, both disease mutant KIs have substantially reduced tubulin transcript levels (Figure S10D) while still displaying no quantifiable differences in tubulin protein levels using Western blot-based detection (Figure S10E); notably, reintroduction of mNG-tagged TTC5 was sufficient to either fully (A231V) or partially (G186V) restore tubulin transcript abundance back to WT levels (Figure S10D). Together, these findings connect the neuronal migration defects to clinically relevant disease mutations, providing new insight into the pathogenesis of this TTC5-associated neurodevelopmental disorder.

### TTC5 loss disrupts neuronal migration and dendritic arborization in the mouse cortex

Since TTC5 loss impairs neuronal outgrowth, arborization and motility in i^3^Neurons, we next sought to determine if the phenotypes we observed extend to neurons in vivo, where we can capture the full complexity of the developing brain. To evaluate neuronal morphology, we utilized a recently-developed approach for genetic manipulation and sparse labeling of individual excitatory layer II/III callosal projection neurons (CPNs)^55,56^. These neurons migrate to layer II/III before extending a primary axon and subsequently elaborating interstitial axonal branches ^57^. Due to the implications for ultimate establishment of correct functional connectivity, the axonal arborization of these neurons is highly stereotyped, each forming approximately 7 to 9 axonal branches in ipsilateral layer V while restricting branching within layer IV ^57^. Following *in utero* electroporation (IUE) of miRNAs to induce TTC5 knock-down, we observed a significant decrease in layer V primary axon collateral branches (5.4+/-1.4 versus 7.5 +/-1.5 branches in controls) (Figure 7A, 7B). Loss of TTC5 had no impact on the lack of interstitial axon branching normally observed in layer IV. Co-electroporation of TTC5 KD miRNAs along with a plasmid overexpressing WT human TTC5 to restore TTC5 levels normalized layer V collateral axon branching (6.8+/-1.4 branches), confirming TTC5 dependence (Figure 7A, 7B). Tracing of basal dendrites in the layer II/III neurons (Figure 7C) revealed TTC5-dependent reductions in total dendrite lengths (1829+/-458 μm for WT *versus* 1140.0 +/-420 μm for TTC5 KD) and branch number (22 +/-7 for WT *versus* 15 +/-6 branches for TTC5 KD), both rescued upon reintroduction of WT human TTC5 (Figure 7D). These in vivo dendritic phenotypes are consistent with the reduced neurite lengths and numbers we observed in the TTC5 KD (Figure 2A, 2B) and R147A KI (Figure 4G) in our 2D i^3^Neuron-glia co-culture paradigm (Figure 2A, 2B).

**Figure 7.**
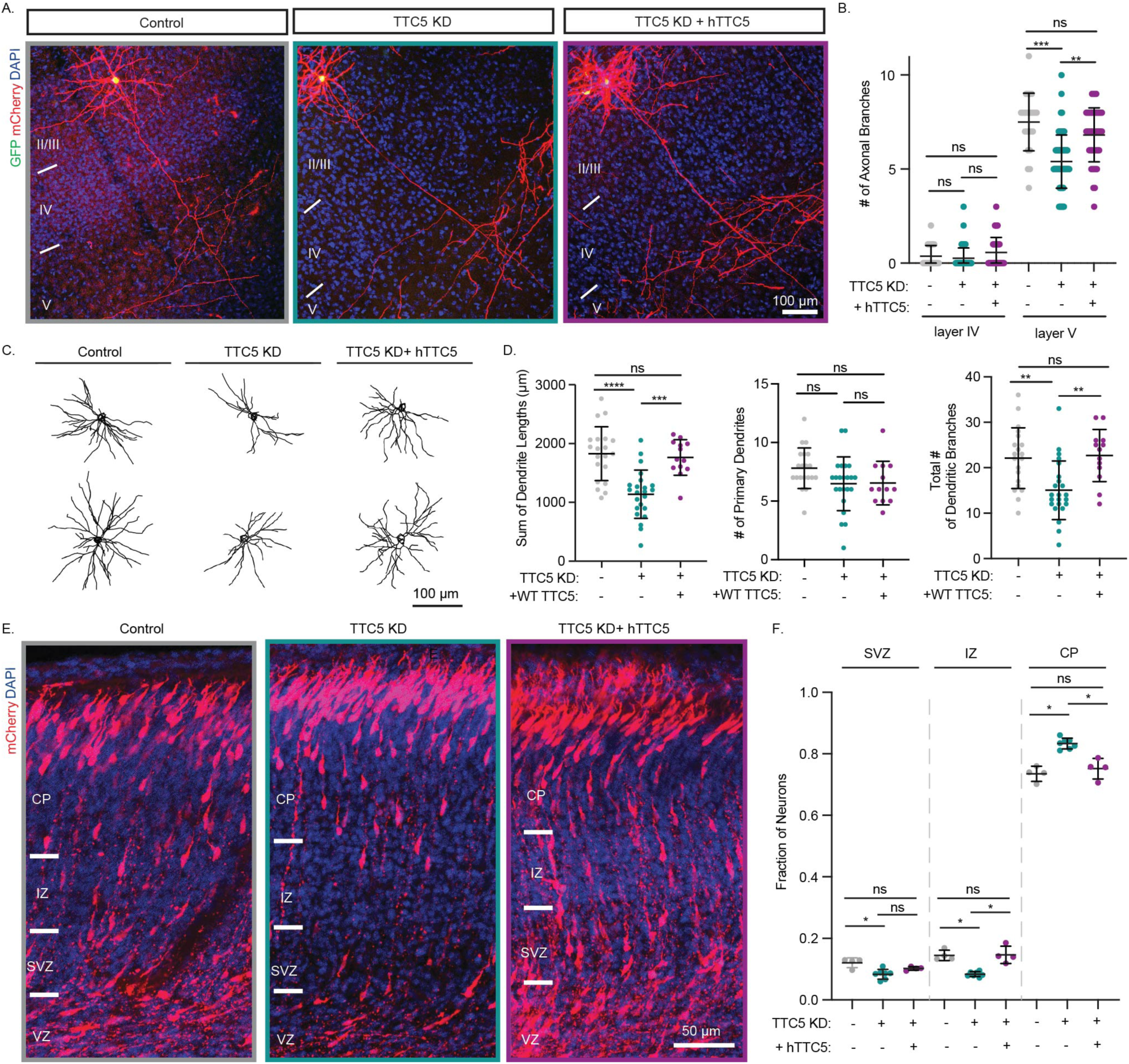
TTC5 KD results in axonal branching, dendritic arborization and migration defects in the mouse cortex that are rescued upon reintroduction of human TTC5. (**A**) Representative images of layer II/III CPNs upon TTC5 KD using miRNA IUE with or without co-electroporation of a human WT TTC5 construct. GFP indicates shRNA expression, mCherry labels neuronal morphology, and DAPI is a nuclear stain for cortical layers delineation. Scale bar: 100 μm. (**B**) Primary interstitial axon branches quantified in layers IV and V for indicated numbers of neurons across multiple mice (control (gray): 24 neurons/7 animals, TTC5 KD (teal): 85 neurons/9 animals, +hTTC5 (purple): 44 neurons/4 animals); bars, mean +/-SD; ns, > 0.05, **, p < 0.01, ***, p < 0.001 using nested one-way ANOVA with Tukey’s correction for multiple comparisons. (**C**) Representative layer II/III dendritic traces from neurons in (A) and quantified in (D). Scale bar: 100 μm. (**D**) Arborization metrics: sum dendrite length, primary dendrite number and total dendrite number (control: 20 neurons/7 animals, TTC5 KD: 23 neurons/8 animals, +hTTC5:13 neurons/4 animals); bars indicate mean +/-SD; ns, p > 0.05, **, p < 0.01, ***, p < 0.001, ****, p < 0.0001 using one-way ANOVA with Tukey’s correction for multiple comparisons. (**E**) Representative images of migrating excitatory neurons (labeled with mCherry and DAPI) in the cortical column at E18.5 upon TTC5 KD or WT hTTC5 co-addition. Scale bar: 50 μm. (**F**) Fraction of total neurons per image distributed across the subventricular (SVZ), intermediate zone (IZ) and cortical plate (CP) (control: 4, TTC5 KD: 7, KD+hTTC5:4 animals); Error bars indicate mean +/-SD, ns, p > 0.05, *, p < 0.05 using one-way ANOVA with Tukey’s correction for multiple comparisons.

We next assessed the impact of TTC5 loss on neuronal migration. Classical migration assays capture gross neuronal mislocalization in the cortical wall, as seen in lissencephaly ^58^. We hypothesized that a postnatal timepoint might be too late to capture a hypermotility phenotype, such as that observed in our TTC5 KD i^3^Neurons (Figure 3). We therefore electroporated E14.5 embryos and analyzed the early migration of neurons from the subventricular zone (SVZ) through the intermediate zone (IZ) to the cortical plate (CP) four days later, at E18.5 (Figure 7E). TTC5 KD resulted in a significant shift in cortical layer distribution, with a greater fraction of neurons reaching the cortical plate (∼83% *versus* 73% in control) and a corresponding decrease in neurons remaining in the SVZ or IZ (Figure 7F). Reintroduction of WT TTC5 rescued this phenotype to control levels (Figure 7E, 7F). Together, these findings corroborate in vivo a pivotal role for TTC5 in both neuron migration and arborization within the higher complexity of the developing mouse cortex, demonstrate the robustness of TTC5-dependent phenotypes across in vitro and in vivo models, and link these cellular defects to the disease etiology of TTC5-associated intellectual disability.

## Discussion

The centrality of microtubule homeostasis to neuronal development and maintenance is evidenced by the breadth of neurodevelopmental and neurodegenerative disorders arising from mutations in microtubule cytoskeletal components and their regulators^21,22^. Motivated by the identification of TTC5 as a key tubulin autoregulation mediator, and its emerging association with neurodevelopmental disorders, we investigated TTC5 and tubulin autoregulation in human neurons. Using iPSC-derived cortical-like glutamatergic neurons, we demonstrate for the first time that tubulin autoregulation is operative in neurons (Figure 1B, 1D), and show through transcriptome-wide profiling, that it dominates the neuronal response to acute microtubule depolymerization with remarkable specificity in both soma and neurites (Figure 1D). While autoregulation was presumed to be universal across tissues, most evidence comes from immortalized cell lines and a limited number of primary cell types ^23,59^. The dramatically attenuated autoregulation we observe at the iPSC stage may reflect lower expression of pathway mediators (Figure S1A), reduced tubulin translating-ribosomes, or allude to the existence of alternate regulatory mechanisms in pluripotent cells.

Although tubulin autoregulation was discovered over four decades ago, the physiological contexts in which it operates have remained elusive. We find that TTC5 loss impairs multiple aspects of early neurodevelopment: neurite arborization, axonal outgrowth (Figure 2) and neuronal motility (Figure 3). That each of these phenotypes is recapitulated by the autoregulation-defective R147A knock-in (Figure 4), by SCAPER knockdown (Figure 5) and rescued by overexpressed wild-type TTC5 but not R147A reintroduction, collectively argues that loss of tubulin autoregulation and microtubule homeostasis underlies these defects – though we cannot fully rule out contributions from an alternate or hitherto unknown function of TTC5. Unexpectedly, TTC5 loss reduces rather than increases microtubule dynamics, counter to the naïve prediction that relieving autoregulatory feedback would increase the soluble tubulin pool and enhance polymerization. A recent study in *C. elegans* similarly found that loss of TTC5 and SCAPER reduces microtubule density in neurons.

It attributed this to a tubulin autoregulation-independent role of constitutively interacting TTC5 and SCAPER in γ-tubulin stabilization and non-centrosomal microtubule nucleation^60^, although tubulin autoregulation has not been directly assessed in that system. This contrasts with the mammalian model, in which proximity-proteomic data indicates that the TTC5-SCAPER interaction is triggered by increased soluble tubulin^12,13^. Our RNA-seq and qPCR data did reveal TUBG1 responsiveness to colchicine-induced autoregulation, hinting at a crosstalk between autoregulation and γ-tubulin-dependent microtubule nucleation. Together, these findings may reflect complementary roles for TTC5 and SCAPER across neuronal maturation with early centrosomal arrays driving polarization, outgrowth and migration, and later non-centrosomal arrays supporting dendrite elaboration and synapse formation ^61^, and with the relative contributions of autoregulation-dependent and -independent activities potentially shaped by the greater complexity of the mammalian nervous system.

The hypermotility phenotype observed upon TTC5 loss (Figure 3) is particularly striking and unanticipated. That it manifests cell autonomously, observed in sparse TTC5 KD neurons cultured within wild-type neurospheres in vitro or in sparse TTC5 KD neurons within the cell-dense environment of the developing cortical column in vivo, argues against indirect or collective migration effects and points to an intrinsic change in motile capacity. While decreased arborization could itself promote directed motility by reducing competing pulling forces on the soma as observed for DCX^52^, the extension of the motility phenotype to non-neuronal HEK293 cells, implies a more universal mechanism. Phylogenetic co-evolution analysis has identified RhoA, RhoB and CDC42, GTPases that are well known regulators of neuronal migration and neurite extension, as predicted functional interactors for TTC5^62,63^. Local microtubule depolymerization releases Rho GTPase activating factors like GEFH1, linking microtubule dynamics to focal adhesion formation and dynamics ^64^. Consistent with this, a recent preprint shows that TTC5 loss in HeLa cells reduces surface expression of adhesion molecules, including E-cadherins, reducing cell-substrate adhesion^51^. We propose that TTC5 loss, by perturbing microtubule homeostasis, dysregulates adhesion pathways to drive changes in motility across different cell types.

Critically, our study provides the first insights into the etiology of TTC5-associated intellectual disability. Patient missense mutations destabilize the TTC5 protein (Figure 6), resulting in TTC5 loss-of-function, and neurons carrying TTC5 disease mutations recapitulate the hypermotility phenotype in our glia-guided migration assay. The in vivo correlate of this – premature arrival of neurons to the cortical plate, with a corresponding depletion from the SVZ and IZ (Figure 7), provides an explanation for the clinical manifestations of cortex thickening and abnormal simplified gyri defects seen in patients with TTC5 mutations ^15^. More broadly, the concordance between iPSC-derived neuronal phenotypes and in vivo findings validates the i^3^Neuron model, with its genetic tractability and accessible differentiation timeline, as a powerful platform for capturing neurodevelopmentally-relevant biology ^65^. In addition, the i^3^Neurosphere glia-guided migration assay we introduce here offers a broadly applicable and scalable readout of neuronal migration, dysregulation of which characterizes many neurodevelopmental disorders ^52,66–70^, and will be amenable for future therapeutic screens.

## Acknowledgements

We thank J. Spector (National Institute of Neurological Disorders and Stroke, NINDS) for imaging assistance and image analysis pipelines, E. Zehr (NINDS) for structure figure panels, V. Ryan, M. Fernandopulle, J. Colon-Mercado, and M.E. Ward (NINDS) for expertise with i^3^Neurons, access to equipment and reagents, S. Anderson and M. Kirby from the Flow Cytometry core (National Human Genome Research Institute) for cell sorting, the iPSC core (National Heart, Lung and Blood Institute, NHLBI) for CRISPR KI lines, the sequencing core (NHLBI) for RNA-sequencing and R. Hogg (NHLBI) for RNA-seq analysis help. We also thank Dr. R. Hegde (Medical Research Council, UK) for HEK-293 TTC5 KOs. This project used computational resources from the NIH High-Performance Computing Biowulf cluster. S.L.S. was partially supported by the PRAT fellowship from the National Institute of General Medical Sciences (NIGMS) Fi2 funding #1FI2GM146657-01. A.R.M is supported by the Intramural Research Programs of NINDS and the NHLBI of the National Institutes of Health (NIH). The contributions of the NIH authors were made as part of their official duties as NIH federal employees, are in compliance with agency policy requirements, and are considered Works of the United States Government. However, the findings and conclusions presented in this paper are those of the authors and do not necessarily reflect the views of the NIH or the U.S. Department of Health and Human. J.Z. is a K99/R00 NINDS postdoctoral fellow (grant number K99NS138586) and A.L.K received funding from NINDS (R03-TR004616) and Johns Hopkins School of Medicine Institutional Funds.

## Methods

### Mammalian cell culture and cell line generation

WT and TTC5 KO HEK293s obtained courtesy of the Hegde Lab^8^ (migration experiments) and HEK293 derivative Lenti-X 293T (production of lentiviruses) cells were cultured in DMEM + Glutamax (ThermoFisher 10564029) supplemented with 10% FBS (Life Technologies A5209402) at 37 °C and 5% CO2. EB1-mNG knock-in U2OS cells were generated as previously described^71^ and maintained in in DMEM + Glutamax (ThermoFisher 10564029) supplemented with 10% FBS.

All induced pluripotent stem cell (iPSC) lines in this study derive from the parental WTC11 line^72^ further engineered to contain 1) a single copy of doxycycline-inducible mouse NGN2 at the AAVS1 locus as previously generated and described ^26,27^ and 2) a single-copy of dCas9-BFP-KRAB within the human CLYBL intragenic safe harbor locus and subsequently termed CRISPRi-i^3^Neuron or i11W-mNC as previously generated and described^28^. iPSCs and i^3^Neuron lines were maintained as described in Fernandopulle et al. 2018 with any deviations to this protocol noted below. Briefly, iPSCs were maintained on dishes freshly coated with Matrigel (Corning 354277) in Essential 8 Medium (E8) (Life Technologies A1517001) or StemFlex (Life Technologies A3349401) supplemented with chroman-1 (C1) (MedChem Express HY-15392) at low density and/or upon passaging and then E8 or StemFlex (SF) alone for subsequent daily media changes. Passaging of iPSCs was performed by washing cells once with PBS followed by 10 minute incubation at 37 ° in StemPro Accutase Cell Dissociation Reagent (A1110501) which upon suspension was diluted 1:1 into PBS containing a small volume of media, centrifuged at 300xg for 5 minutes and resuspended into either SF for replating or N2 induction media for subsequent differentiation. All stocks of iPSC cell lines were prepared by freezing into KnockOut Serum Replacement (Life Technologies 10828028) supplemented with 10% DMSO and stored in liquid nitrogen. Throughout, unless noted, differentiations into neural progenitor cells (NPCs) and i^3^Neurons were carried out on Matrigel-coated dishes via addition of an N2 induction media composed of knockout DMEM/F12 medium (Life Technologies 12660012) with addition of N2 supplement (Life Technologies 17602048), 1xNEAA (Thermofisher 11140050), 1xGlutaMAX (Thermofisher 35050061), doxycyline (Takara 631311) and chroman-1 (C1)) on days 0, 1 and 2 (with PBS washes as needed) followed by replating of the D3 pre-differentiated neurons. Recipient plates were freshly coated for at least 2 hrs and up to overnight incubation at 37 ° with poly-L-ornithine (PLO) (Sigma P3655). PLO was prepared fresh at 0.1 mg/ml in 10% borate buffer (pH 8.4) (A09-2053-100, Aniara Diagnostica Inc.) in cell culture grade water (25-055-CV, Corning) and filtered before addition to plates. Following incubation, plates were washed four times with water and fully dried before addition of cells. When stated, PLO-coated plates were subsequently coated with laminin (R&D Systems #3446-005-01) resuspended in cold PBS (at 15 μg/ml) and incubated at 37°C for 1.5 hours. Cells were then resuspended in Accutase, washed, spun and resuspended in the i^3^Neuron culturing media referred to as “BP++” composed of BrainPhys medium (Stemcell Technologies 05790) with 1xB27 supplement (ThermoFisher A3582801), BDNF (PeproTech, 450-02), NT-3 (PeproTech 450-03), Cultrex 3D Culture Matrix Laminin I (BioTechne, 3446-005-01) and doxycyline (2 μg/ml) as previously described. Laminin was removed from the recipient plate, followed immediately by plating of cell suspension or addition of BP++ for subsequent plating of neurospheres (described in detail below).

The reporter cell line expressing cell-fill mScarlet and EB1-EGFP used for all morphological (axonal outgrowth, arborization) and microtubule dynamics readouts were generated through lentiviral delivery of pLenti-EB1-EGFP (Addgene: 118084 pLenti-EB1-EGFP was a gift from Ken-Ichi Takemaru (Addgene plasmid # 118084; http://n2t.net/addgene:118084; RRID:Addgene_118084) to the i2 mScarlet-expressing CRISPRi-i3N iPSC line generated and described in Tian et al. Individual subclones were picked by manual scraping using the Lumascope (EtaLuma) from distinct colonies grown up following replating of 1000 cells onto a 10 cm dish and screened for co-expression of red and green fluorescence. A single subclone was selected for all downstream experiments to ensure uniform expression of EB1-EGFP and mScarlet between experimental conditions. A normal karyotype for the clonal line was confirmed by G-banded karyotyping performed at passage 3 post-transduction by WiCell Cytogenetics lab. (Madison, WI) using twenty randomly selected metaphases. This reporter line then served as the parental line for subsequent sgRNA knock-down of our genes-of-interest and generation of Halo-NLS+ lines via lentiviral transduction for subsequent monitoring of neuronal nuclei (see neurosphere migration assay). The Halo-NLS construct was obtained courtesy of the Ward Lab. The mScarlet-fill/EB1-EGFP reporter line also served as the parental line for subsequent knock-in of TTC5 point mutants R147A, A231V and G186V performed and validated by the NHLBI iPSC core. Briefly 1×10^6 iPSCs were transfected with premixed 20 μg HiFiCas9 V3 (IDT#1081061), 200 pmol sgRNA (Synthego) (R147A: CAATGGTGCTTCGTCAGCTG, A231V: CCTTCATCTGAACAGGGCGA, G186V: TTTCTCTTTCAGATATTCTT) and 200 pmol ssODN (IDT) (R147A: agTGCAGGAACAAAGTCTCCTTGCAAAACCTGTCAATGGTGCTTGCTCAGCTGCGGACTGACACTGAAGATGAACATTCTCACCATGTCATGGACAGTGT, A231V: GAAAGTTGACAGAAAAGCTTCTAGCAATCCTGACCTTCATCTGAACAGGGTGACGgtaagacctttggggtgtagtcatctgagggtcaaataaataaga, G186V: ttgaacttggactctgagatgatttgtcactgtgtttctctttcagATATTCTTGTGAATTCATATCTTTCCCTTTACTTCTCTACTGGCCAGAACCCTA) using Nucleofector 4D with buffer P3 and program CA-137 (Lonza) and then plated onto one well of rhLamin521 (Thermo) coated 6-well plate with StemFlex, RevitaCell (Thermo) and HDR enhancer V2 (IDT). Transfected cells were cultured in 32°C incubator for 3 days before moving to 37°C incubator, and fresh medium was changed daily after transfection. Single-cell subcloning was done using manual serial dilution to 1 cell/96-well Matrigel coated plate with StemFlex and CloneR2 (Stem Cell Technologies). 10 days later, single-cell clones were picked and confirmed by genomic PCR (R147A Forward primer: gcaagactccacctcaaaca; Reverse primer: gcccttccttgcttacgtg, A231V Forward primer: GGGACTAAGATATCCACACTGTTAT; Reverse primer: CCCCATTCTCTACCTATTCTTCCA, G186V Forward primer: GCCAGAGAGGGCATCTTTGTT; Reverse primer: TGATGCTTGCTAAAGGGCCA), Sanger sequencing, and ICE analysis (Synthego). An unedited clone that had undergone the full parallel process of clone selection served as the WT comparison for the selected R147A and A231V or G186V knock-in clonal line and when possible multiple knock-in clones were tested in each assay to assess and rule-out clone-to-clone variability. Disease mutant rescue knock-in lines expressing WT TTC5-mNeonGreen (mNG) and Halo-NLS were generated via lentiviral transduction of TTC5-mNG-pLVX-TetONE-Puro and Halo-NLS constructs into the indicated knock-in clone followed by puromycin selection. The tagged TTC5 construct was generated via NEB HiFi cloning of human WT TTC5 cDNA obtained from Genscript (OHu06087) and mNeonGreen fragment into backbone pLV-TetOne-Puro-hAXL, a gift from Kenneth Pienta (Addgene plasmid # 124797 ; http://n2t.net/addgene:124797&tab;; RRID:Addgene_124797).

Rescue experiments in the TTC5 KD were performed by co-transducing non-fluorescent NT control and TTC5 knock-down lines with either a WT or R147A mutant TTC5-mNG construct and the Halo-NLS construct. The distribution of TTC5-mNG fluorescence signal was similar for both WT and R147A constructs as assessed by flow cytometry and cells were subsequently sorted for medium-high mNG+ and Halo-NLS+ expressors. The TTC5-mNG fragment was cloned out of the TTC5-mNG-pLVX-TetONE-Puro vector (above) and inserted into backbone pCDH-EF1a-eFFly-mCherry, a gift from Irmela Jeremias (Addgene plasmid #104833 ; http://n2t.net/addgene:104833 ; RRID: Addgene_104833). The R147A mutation was then introduced into this construct via site-directed mutagenesis. NT and TTC5 KD lines expressing mNG alone (not attached to TTC5) were generated in parallel via co-transduction of “J81” (EF1a-mNeonGreen plasmid courtesy of the Ward Lab) and the Halo-NLS construct and sorted for co-expressors as above (of note, mNG fluorescent signal was much higher for the mNG-only construct relative to the mNG-tagged TTC5 constructs).

Glia were obtained commercially as mouse C57 Brain Mixed Astrocytes (M-ASM-330 Lonza), expanded and cultured in “glia media” AGM Astrocyte Growth Medium (CC-3186 Lonza) prior to co-culture with i^3^Neurons in BP++. To prepare glia for co-culture, glia were added in a 25 ul drop (at a density of approximately 50,000 cells per well) to the center of a matrigel-coated and air-dried Ibidi and topped off after 10 mins at 37° with 325 ul glia media. A full glia media change was performed the following day and cells maintained with ½ media changes every 3-4 days prior to replating of D3 i^3^Neurons in BP++ media.

### Drug treatments

Microtubule depolymerization was performed by administration of colchicine (Sigma C9754) resuspended in H20 as a 100 mM stock and diluted to a final concentration of 10 μM in corresponding media.

### Lentivirus Production

For preparation of lentiviruses, 2.5 million pLentiX cells were seeded in each well of a 6-well plate precoated with PLO (incubated at 37° 1 hr, washed 4 times with H20 and fully dried). Typically, three wells of a 6-well plate were pooled per sgRNA construct. The following day, cells were changed to fresh media and transfected with the lentiviral vector (1.2 ug/well) and a mastermix of three plasmids psPAX2, pMD2G, and pAdVantage (at a ratio of 1.6 ug: 0.6 ug: 0.2 μg respectively) using Lipofectamine 3000 (ThermoFisher L3000001) according to manufacturer’s instructions. At 18 hours post-transfection, cells were washed once with PBS and replenished with fresh DMEM (3 mls/well) supplemented with viral boost (Alstem VB100) at 1:500 dilution. Forty-eight hours later, virus-containing media was pooled per construct upon collection into 15 ml conicals and spun at 1100xg for 5 mins, filtered through a 0.45 μm PES filter into a fresh 15 ml conical followed by addition of Lenti-X Concentrator (Takara 631232) at a ratio of 1:3 concentrator to viral supernatant and incubated at 4 ° for 48 hours. After 48 hours, virus was concentrated by centrifugation at 1500xg for 45 mins at 4°, supernatant removed and viral pellet resuspended in 0.1X the original collection volume in cold PBS and frozen in single-use aliquots at -80°.

### sgRNA Cloning, Packaging and Transduction of iPSCs for CRISPRi knock-down line generation

sgRNAs oligos targeting the promoter of each studied gene were selected from the hCRISPRi version 2 (v2) library designed with overhangs for ligation into the sgRNA expression vector ‘pCRISPRia-v2’ and cloned as described in Horlbeck et al. 2016 and packaged into lentiviruses as described above ^73^. The sgRNA oligos used in this study are as follows:

B9 non-targeting control: F: TTAGCTCTTAAACTAGGTGCTTGCGCTTAGTCCCAACAAG, R: TTGGGACTAAGCGCAAGCACCTAGTTTAAGAGC

TTC5 sg1: F: TTAGCTCTTAAACACCTGACTGCGGTTAATTTCCAACAAG, R: TTGGAAATTAACCGCAGTCAGGTGTTTAAGAGC

TTC5 sg5: F: TTAGCTCTTAAACACGAGGCTCCCTGGCTGGCCCAACAAG, R: TTGGGCCAGCCAGGGAGCCTCGTGTTTAAGAGC

CNOT10 sg1: F: TTAGCTCTTAAACACTACTTCCGGGGGCGGCACCAACAAG, R: TTGGTGCCGCCCCCGGAAGTAGTGTTTAAGAGC

CNOT11 sg2: F: TTAGCTCTTAAACCCGAGTGTGGCCTGGGCTACCAACAAG, R: TTGGTAGCCCAGGCCACACTCGGGTTTAAGAGC

SCAPER sg1: F: TTAGCTCTTAAACCCCTTCGTTTGACAGGTGACCAACAAG, R: TTGGTCACCTGTCAAACGAAGGGGTTTAAGAGC

For all CRISPRi knock-down experiments, 8.3×10^4 iPSCs were transduced with 50-100 μl of concentrated lentivirus stocks, washed the following day with 2x PBS and replenished with fresh E8+C1 for recovery, followed by rapid selection at 48 hours post-transduction with puromycin (10 μg/ml). Upon reaching confluence, transduced cells were utilized in assays or frozen down as low passage post-transduction stocks.

### siRNA knock-down

For siRNA knockdowns, 3.75×10^4 U2OS cells were reverse-transfected with Silencer Select siRNAs (reconstituted in H20 and stored at 20 μM stock solution) using Lipofectamine RNAiMAX to a final concentration of 40 nM according to manufacturer instructions. siRNA Silencer siRNAs (ThermoFisher) used in this study include Negative Control No. 1 (4390843) and TTC5 (s40806).

### RNA extraction, cDNA preparation and RT-qPCR

All RNA extractions were carried out using the Zymo Direct-zol RNA Mini- (R2053) or MicroPrep (R2063) Kits according to manufacturer’s instructions and quantified by nanodrop for generation of cDNA from 500-1000 ng of RNA input using the High-Capacity cDNA Reverse Transcription Kit (Life Technologies 4368814). RT-qPCR was performed using either Powerup Sybrgreen MasterMix (ThermoFisher A25777) with the primer pairs either previously described as in Lin et al^8^ or designed using NCBI PrimerBlast (TTC5-F:GGCTTCACCGAATTCAGCAC/R:TGAGGCTTCCCATTCACCAC, SCAPER-F:TTCCAGCGCTCCAATAGTCA/R:TTCCGAGGGTGCCTTGTTTT, CNOT10-F:CGTCTGCCCACTCCTCTAGC/ACTGACCTGTGCCTTCATGTT, CNOT11-F:TTCAAAAAGACGCCTCGCCA/R:GAGGCGGTGGACGAATAAAC, TUBB-F:GAAGCCACAGGTGGCAAATA/R:CGTACCACATCCAGGACAGA, TUBA1A- F: CCACAGTCATTGATGAAGTTCG/R:GCTGTGGAAAACCAAGAAGC TUBG1- F:GCTCTGGACTGGGTTCCTAC/R: ATGGGTTCTGGATGTGCAGG, GAPDH-F:AACATCATCCCTGCCTCTACTGG/GTTTTTCTAGACGGCAGGTCAGG) or Taqman Advanced Fast MasterMix (ThermoFisher 4444963) using commercially-available probe/primer sets (ThermoFisher; TUBB [Hs00962420], TUBB3 [Hs00964962], TUBA1A [Hs03045184], TTC5 [Hs00381484] and HPRT [Hs02800695]) for i^3^Neuron RNA quantification and run on a QuantStudio 6 Flex Real-time PCR System according to manufacturer’s instructions. Each sample was assayed in technical pipetting duplicate. For analysis, cDNA input was normalized (dCt) to GAPDH or HPRT respectively and expression fold changes were calculated using the ddCt method to a normalizing control (ex. the untreated condition).

### RNA-Seq: cDNA Library Preparation, Sequencing and Analysis

For separation of proximal/distal neurite and soma/neurite (i.e. total cell) RNA fractions for subsequent RNA sequencing, 1 μm pore size cell culture inserts (Falcon 353102) as previously described ^74,75^ (and validated for the i^3^Neuron model^76^) were added to 6-well plates followed by sequential PLO and laminin coating, seeded with 3×10^6 D3 pre-differentiated cells per insert and maintained with half-media changes every other day for the duration of the experiment (ie. 1 mL of BP++ removed from top and bottom and 2 mLs of fresh BP++ added to the top). At 14 days post-differentiation, cells were either left untreated or treated for 5 hours with colchicine, followed by separate harvesting of the neurite and soma/neurite fractions by cell scraper. The separate wells were pooled per condition to obtain sufficient material from the neurite fraction, pelleted and resuspended in TriReagent (Zymo) for subsequent RNA extraction (by Micro and MiniPrep Kits respectively) and quantification using the Quant-IT RNA Assay Kit (Life Technologies Q33140) according to manufacturer’s instructions. Sequencing libraries of the soma/neurite fraction were prepared from 40 ng input using the QuantSeq-Pool Sample-Barcoded 3’ mRNA-Seq Library Prep Kit (Lexogen) according to manufacturer’s instructions and sequenced by the NHLBI sequencing core using the Illumina NovaSeq 6000. For analysis, raw fastq reads were trimmed using the fastp package ^77^ (with parameters --detect_adapter_for_pe --trim_poly_g) and aligned to the hg38 genome using Pipeliner, an open-source analysis pipeline (https://github.com/CCBR/Pipeliner) run through the NIH high-computing Biowulf cluster. Differential expression analysis was performed using Voom/Limma through the Degust server ^78^ (Powell DR (2015) Degust: interactive RNA-seq analysis. https://doi.org/10.5281/zenodo.3258933) with a cut-off of minimum gene read count of 20 and a minimum CPM >5 in at least 1 sample. Raw data and sequencing files are deposited with the NCBI Gene Expression Omnibus (GEO) database with accession number GSE312628.

### Western blot analysis

For preparation of whole-cell lysate, cells were washed with PBS and lysed in RIPA buffer (50 mM Tris-HCl, 150 mM NaCl, 0.1% SDS, 0.5% NaDoc, 1% TritonX-100 pH 7.5) with addition of cOmplete™ Mini EDTA-free Protease Inhibitor Cocktail Tablets (Sigma), incubated on ice for 15 minutes with intermittent vortexing and clarified at 14000 x g for 20 minutes at 4° C. Loading samples were prepared with 4x sample loading dye ([106 mM Tris HC, 141 mM Tris Base, 2% LDS, 10% glycerol, 0.51 mM EDTA, 1% Coomasie Blue, 0.175 mM Phenol Red pH 8.5] and 0.1 M DTT) and run on a NuPAGE 4-12% Bis-Tris Gel in either MES (50 mM MES, 50 mM Tris base, 0.1% SDS, 1 mM EDTA, pH 7.3) or MOPS (NuPage 20xSDS MOPS buffer diluted to 1x in sterile MilliQ water [Thermo Fisher Scientific NP0001]) and transferred to a nitrocellulose membrane (LifeTechnologies 88025) using the Trans-Blot Turbo Transfer System (BioRad) or Genscript eBlot L1. The membrane was blocked in 5% milk in PBST for 1 hour and then incubated with primary antibodies overnight at 4°, followed by 3 washes with PBST and a 1 hour incubation at room temperature with secondary antibodies (Licor IRDyes 680 and 800CW goat-anti-mouse and goat anti-rabbit igG (H+L) LIC-926-68020, LIC-926-32211). The membrane was washed 3 times with PBST followed by a final wash with PBS and imaged on the Odyssey CLx system. Primary antibodies used in this study include: anti-TTC5 (NBP1-76636, Novus Biologicals or Abcam AB36855), anti-CNOT10 (15938-1-AP, VWR), anti-CNOT11 (SC-377068, Santa Cruz) and anti-SCAPER (Abnova; H00049855-B02P) for validation of sgRNA knock-downs, and mouse-anti-GAPDH (D4C6R-97166, Cell Signaling Technologies) or rabbit-anti-GAPDH (14C10-#2118, Cell Signaling Technologies) for normalization. Tubulin levels were assessed using antibodies to α-tubulin (DM1a; T9026, Sigma) and TUBB3 (801201, BioLegend).

### i^3^Neurosphere assay for quantification of axonal outgrowth and microtubule dynamics

To prepare “neurospheres” or clusters of differentiating iPSCs in suspension for subsequent replating and outgrowth assays, 10,000 iPSCs resuspended in 30 μl of N2 induction media were added to each 96-well of a low attachment spheroid microplate (Corning CLS4520) that had been pre-incubated with 50 μl N2 per well. Upon addition, plates were spun at 500xrcf for 10 minutes to center cells and left overnight at 37°. The following day an additional 120 μl of N2 induction media was added. Alternatively, spheres could be prepared in 384-well low-adherence format (Corning 12456722) following the same protocol but with addition of 60 ul of N2 the day after preparation. On D3 post-induction, media was replaced with BP++, neurospheres were collected into a wide-orifice p200 pipette tip and replated into PLO/laminin-coated 8-well Ibidi dishes (Ibidi, 80827-90) or 24-well glass bottom dishes (Cellvis, P24-1.5H-N) containing 300 ul BP++. Half media changes were performed every other day for duration of experiment coordinated to fall immediately following EB1-EGFP comet imaging sessions. While individual neurospheres therefore technically constitute individual “differentiations,” for certain experiments an entirely separate neurosphere generating session was often included as another layer of biological replicate for phenotype validation/robustness as stated in the accompanying figure legends.

All confocal images in this study were acquired using a Nikon Ti2 with CSU-W1 spinning disk confocal microscope outfitted with Photometrics Prime BSI or Hamamatsu Fusion BT camera and Nikon LU-NG laser system with 405 nm, 488 nm, 561 nm and 647 nm solid-state diode lasers. To monitor axonal outgrowth, neurospheres were imaged at D4, D6 and D8 for cytoplasmic mScarlet using a 10x/0.3 N.A. objective (Nikon, Plan Fluor 10x OFN25 DIC N1) as 30 μm Z-stacks (step size: 6 μm) tiled as needed to encompass outgrowth over the course of the experiment. NIS Elements AR Analysis microscope imaging software (5.21.03) was then used to quantify the radius of the neurospheres as a readout of axonal outgrowth. Briefly, max-projected images were thresholded on either the high intensity inner sphere of cell bodies or to encompass the lower intensity outer neurites and “equivalent diameter” (the diameter of a sphere corresponding to the thresholded area) calculated for each. The radius of the neurites was then normalized by subtracting the radius of the inner sphere to yield the “axonal outgrowth radius” as previously described^44^. For rescue experiments in the TTC5 KD, neurospheres were instead imaged for mNG signal corresponding to the tagged WT or R147A TTC5 or mNG only protein. As this signal was weaker than the mScarlet typically used for these experiments, max-projected images were manually traced as ROIs around the neurites while still performing automated thresholding on the inner sphere. These ROIs were then used to calculate the “equivalent diameter” as above. For quantification of microtubule dynamics in growth cones, videos of EB1-EGFP (30 frames at 2 sec interval for a total length of 1 min) encompassing multiple growth cones at the growing edges of the D5 neurosphere were obtained using a 60x water objective (N.A. 1.27). For analysis, images were cropped on individual growth cones taken from multiple fields of view and multiple wells and analyzed using the open source uTrack software (https://github.com/DanuserLab/u-track) in MATLAB according to the provided documentation. The resulting individual EB1 comet velocities were filtered for displacement greater than 2 μm and averaged to yield mean comet velocity per growth cone. Comet densities were calculated by dividing the number of comets per growth cone (with displacement greater than 2 μm) by the area of the growth cone determined by thresholding on background levels of EB1-EGFP fluorescent signal. For NT and TTC5 siRNA knock-down EB1-mNG knock-in U2OS cells, videos were acquired as above for growth cones and analyzed by uTrack (as described above) on a 15 μm ^2^ region of cytoplasm (not encompassing centrosome) taken across multiple FOV and wells per condition.

### Neuronal Arborization Assay in i^3^Neurons

D3 i^3^Neurons expressing cell-fill mScarlet were seeded at low density (175 neurons/well) onto D3 cultured glia (seeded in 8-well Ibidi plates as above) and assayed on D12 for arborization via longitudinal imaging of scarlet fluorescence every ten minutes over eight hours at 10x magnification tiled across the entire well. D12 was selected as the optimal timepoint at which the neuron displays appreciable neurite numbers while minimizing crossover between its own neurites or from the axons of surrounding cells. ROIs encompassing a 300 μm radius around individual soma were then selected from a stitched field of view followed by automated analysis with a custom Nikon GA3 analysis pipeline to quantify metrics of arborization such as sum of neurite lengths (following exclusion of the axon), number of neurite ends and neurite branchpoints used in turn to estimate primary neurite number. For each neuron, metrics were averaged across all frames to account for the continued dynamicity of the neurites at this timepoint. Within each experiment, neurons were selected and pooled across 2-3 separate wells.

### Cell proliferation Assay

Following CRISPRi-mediated knock-down of respective genes or knock-in of the R147A mutation, iPSC lines were transduced (as above) with lentiviruses to deliver a Halo-NLS construct. Upon recovery and expansion, iPSCs were seeded at 2500 cells per well into a 96-well plate that had been coated with PLO, washed four times with water, air-dried and subsequently coated with Matrigel. Later the same day, iPSCs (Day -1) were imaged by spiking in Halo ligand JF646 (Promega; reconstituted to 20 μM stock in DMSO) to a final concentration of 10 nM in E8 +C1. The following day, iPSCs were imaged (Day 0) and differentiation initiated via addition of N2 media. Cells were imaged at 10x magnification on each subsequent day with N2 media changes supplemented with Halo ligand JFX646 out to Day 2, then switched to BP++ media on Day 3. Images were analyzed using a custom Nikon elements pipeline to segregate and count number of nuclei per well and averaged across 10 wells per condition.

### Neuronal migration Assay in i^3^Neurospheres

To promote migration of the neurons out of the central sphere, neurospheres generated (as above) from the Halo-NLS+ reporter lines were plated onto D3 cultured glia instead of PLO/laminin. Spheres were then assayed longitudinally to monitor the nuclear distribution between the sphere and underlying glia. Specifically, spheres received fresh Halo ligand JF646 (10 nM final concentration) in half media changes performed the day prior to each imaging session. Each sphere was then imaged at 10x magnification as 45 μm Z-stacks (step size: 5 μm), tiling as needed to accommodate dispersion over the timecourse of the experiment (D6, D9, D12). Max-projected images were analyzed using a custom ImageJ macro to generate an intensity profile of the nuclear signal across a line extending from the center of the sphere to the periphery and rotated radially to obtain an average across every radian of the sphere. These profiles were then averaged across multiple spheres to generate averaged profiles per condition. For the migration tracking analysis, lower-labeling densities required for tracking of individual nuclei were achieved by generating neurospheres of 7.5% Halo-NLS labeled CRISPR KD or CRISPR KI lines spiked into an unlabeled parental control line. These neurospheres were then plated onto glia, labeled with Halo ligand JF646 on D7, and imaged at 10x (tiling as needed) every 15 minutes for up to 22-24.5 hours from D8-D9. Automated tracking was performed using Trackmate, thresholding on intensity and filtering for individual nuclei based on a size cut-off of a radius between 3 and 7.4 μm. Nuclei across frames were linked using simple LAP tracker (linking max distance=22 μm, gap-closing max distance =45 μm and gap-closing max frame gap=3) and then filtered for tracks with greater than 40 spots (indicating nuclei present and continuously tracked for ∼1/2 of the total imaging time).

### Migration Assay in HEK293s

Halo-NLS+ WT and TTC5 KO HEK293s were generated via lentiviral transduction of the Halo-NLS construct followed by sorting for Halo-NLS+ cells. WT and TTC5 KO reporter HEK293s were then resuspended, singularized through a blue cell strainer-capped Falcon tube (FisherScientific 0877123) and seeded at approximately 6000 cells/well onto glass-bottom 8-well Ibidi plates pre-coated with 40 μg/ml fibronectin (Sigma F2006) (resuspended in PBS) and incubated at 37° for 40 mins followed by two washes with culture media. Cells were allowed to adhere for 6 hours prior to imaging every 15 minutes for a total of 8 hours. Automated tracking was performed using Trackmate, thresholding using LoG detector with an estimated object diameter of 41 μm and filtering on quality threshold > 0.85 and max intensity >∼460. Nuclei across frames were linked using simple LAP tracker (linking max distance=50 μm, gap-closing max distance =30 μm and gap-closing max frame gap=4) and then filtered for tracks with greater than 10 spots (indicating nuclei present and continuously tracked for ∼1/3 of the total imaging time) and track max speed <0.04 μm /sec.

### Animals

Experiments were carried out in strict accordance with the recommendations in the Guide for the Care and Use of Laboratory Animals of the NIH. The animal protocol was approved by the Animal Care and Use Committees of the Johns Hopkins University School of Medicine (Protocol #:MO23M68). Timed pregnant CD1 wild-type females were obtained from Charles River Laboratory. The day of birth was designated as P0. Mice of both sexes were used in all experiments and were housed in a 12:12 light-dark (LD) cycle.

### Plasmids and in Utero Electroporations

Plasmids used for the bipartite labeling method are as described specifically pCAG-Cre-ERT2, pEF1-Flex-FlpO and pCAG-FSF-mCherry ^55^. Supernova plasmids are as described ^79^. Plasmids used for knockdown experiments were prepared by inserting modified microRNA-30 (miR-E) at the 3’ end of EGFP in the pCAG vector ^80^. Sequences for RNA silencing of TTC5 were TTCATCTCCAGAGTCGGTCTGC, TCTCGAAAACAGTAGAGCCGAT and TCAGGAGTCACATTCAGTGCCT. DNA encoding human TTC5 was amplified from a human *TTC5* ORF (Sino Biological, HG23976-U) and either cloned in pCIG-FSF vector ^56^or fused at C-terminus with 3x ALPHA tag and cloned into pCAG vector as pCAG-FSF-TTC5-IRES-GFP while control vector was pCAG-FSF-GFP only. In all cases, multiple fragments were assembled using a HiFi NEBuilder kit. Lack of an available immunofluorescence-compatible antibody to perform TTC5 KD validation motivated validation via phenotypic rescue upon WT TTC5 construct readdition. All new constructs were verified by restriction reaction and sequencing. For *in utero* electroporation, endotoxin-free plasmid DNA was prepared using NucleoBond® Xtra Midi EF kit.

*In utero* electroporations were performed as described ^56^. Briefly, timed pregnant E14.5 (neuron migration experiments) or E15.5 (neuronal morphology analysis) mice were anesthetized with isoflurane and placed on a heating pad. The incision site was shaved, cleaned, and local anesthetic (Bupivacain) was applied subcutaneously. A small laparotomy was performed, and the embryos were removed and rinsed with warm sterile PBS. The lateral ventricles were then injected with small volumes (approximately 0.5–1 µl) of endotoxin-free DNA solutions in PBS with Fast Green dye. Embryos were electroplated with gene paddles (Harvard Apparatus) using a BTX square pulse electroporator (parameters: 40 V, three 50-ms pulses with a 950-ms interval). In each experiment, equal number of control and experimental embryos were electroporated. Animals were placed on a heating pad to recover and pain was further controlled by subcutaneous injection of Carprofen. Following birth, pups were injected with Tamoxifen at P1 and P2 as described ^56^.

### In Vivo Neuronal Migration Assay and Analysis

Embryos electroporated at E14.5 were analyzed at E18.5 to assess position of migrating excitatory neurons in the cortical column. Pregnant mice were sacrificed by cervical dislocation and embryos quickly removed, rinsed in cold PBS and decapitated. Heads were then fixed overnight at 4 °C in 4% PFA dissolved in PHEM buffer (27 mM PIPES, 25 mM HEPES, 5 mM EGTA, 0.47 mM MgCl_2_, pH = 6.9) with 5% sucrose and 0.1% Triton-X100. Then, all samples were washed multiple times in PBS, brains isolated, sliced coronally (150 µm sections) and incubated in a blocking buffer for at least 30 min at the room temperature (3% BSA with 0.3% Triton-X100 and 0.02% Sodium Azide in PBS, filtered). Brain sections were incubated with primary antibodies (chicken-anti-GFP [AVES GFP-1010], rabbit-anti-dsRed [Takara 632496], mouse-anti-ALFA [Nanotag-N1582]) dissolved in a blocking buffer with 5% normal goat serum (NGS) overnight at 4 °C. Slices were then washed three times (each round at least 1 h, room temperature) in PBST (PBS with 0.1% Triton-X100) and incubated in secondary antibodies and DAPI dissolved in PBST overnight at 4 °C. Following another three washes, slices were mounted on a glass slides using a Fluorogel/DABCO. Samples were imaged with inverted Zeiss 700 confocal microscope using a 20× objective (Plan-Apochromat 20×/0.8). Imaging parameters were as follows: px size 0.31 × 0.31 µm (2048 × 2048 px resolution at 0.5 zoom, total image size was 640 × 640 µm), optical sections 1.8 µm, z-stack 1.5 µm. 3 - 5 slices with electroporated cortical regions were captured per animal. Confocal image stacks were manually analyzed in ImageJ. Neurons were counted in the subventricular zone, intermediate zone, and cortical plate, and numbers were then normalized to total number of neurons in the column.

### In Vivo Neuronal Morphology Analysis

Animals were transcardially perfused with PHEM buffer followed by ice-cold 4% PFA/PHEM with 5% sucrose and 0.1% Triton-X100. Brains were isolated and post-fixed overnight at 4 °C. Then, all samples were washed in PBS, sliced coronally (250 µm sections), and processed for immunostaining as described above. Confocal image stacks were manually analyzed in ImageJ with primary collateral axon branches were scored along the principal axon in the layer 4 and layer 5 as previously described ^56^. Dendrites were manually traced in Neurolucida360 Software.

### Statistical Analysis

Statistical analysis was performed as stated in accompanying figure legends using GraphPad Prism 9 and 10. Single, double, triple and quadruple asterisks indicate p<0.05, p<0.01, p< 0.001 and p<0.0001 respectively. For in vivo work, branching data and Migration data were statistically analyzed by fitting a mixed effects model (Maxwell et al, <u>2018</u>) as implemented in GraphPad Prism 9.5.0. This mixed model uses a compound symmetry covariance matrix and is fit using Restricted (Residual) Maximum Likelihood (REML). Significance was determined with either a *t*-test or a one-way ANOVA with a post-hoc Dunnet’s test.

## Supplemental Figures

**Figure S1.**
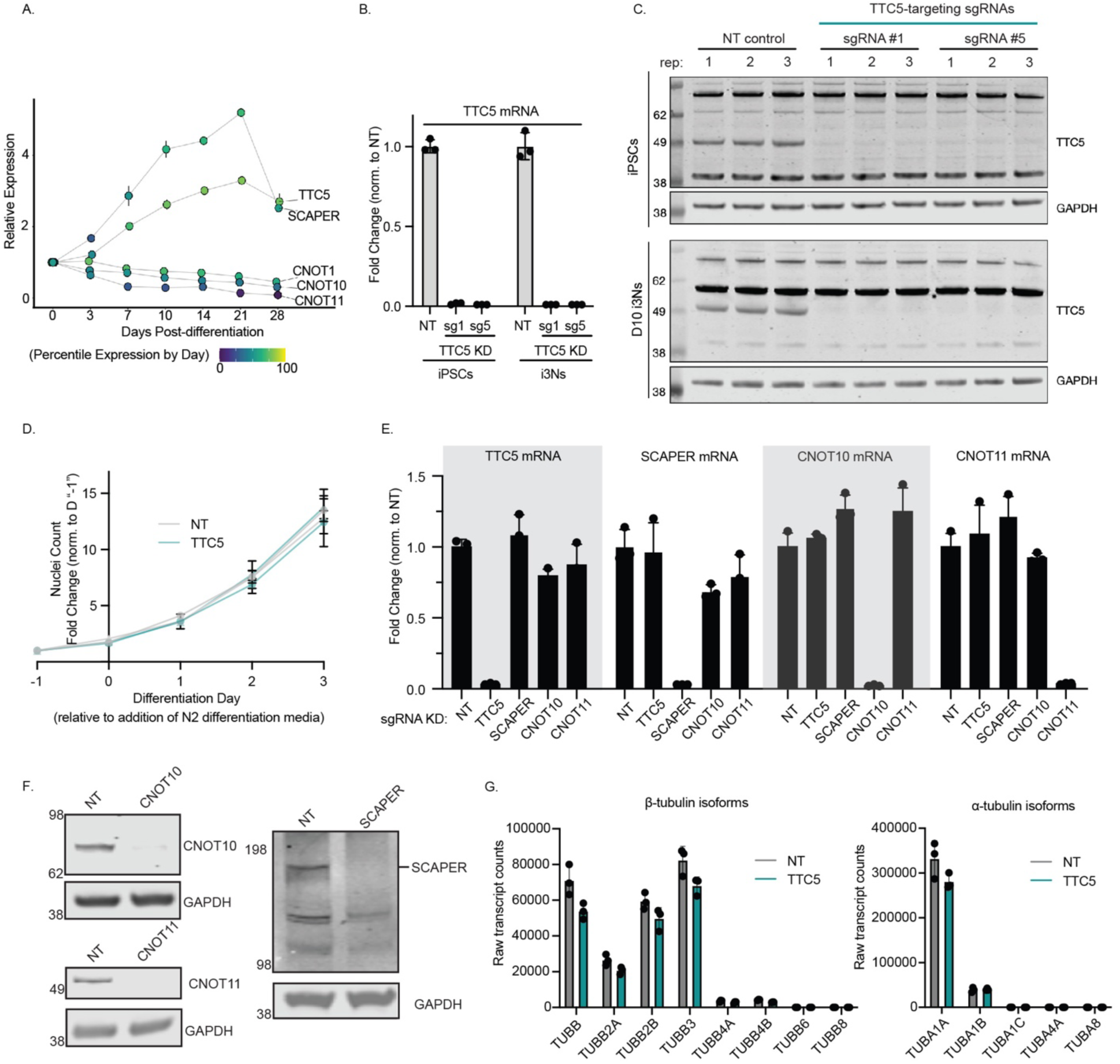
Baseline characterization and validation of tubulin autoregulation effector KDs from iPSCs to i^3^Neurons (related to Figure 1). (**A**) Relative protein expression levels of indicated genes across a timecourse of i3N differentiation^25^ generated using https://niacard.shinyapps.io/i3Neuron/. (**B - C**) Control and TTC5 KD iPSC lines were generated in parallel via lentiviral transduction of iPSCs with two independent TTC5 promoter-targeting guides or a non-targeting (NT) control followed by selection and differentiation into i3Ns. At iPSC stage or D10 post-differentiation, cells were harvested and assayed for TTC5 mRNA and protein expression by (**B**) RT-qPCR, normalized to HPRT and NT control and (**C**) Western blot with GAPDH as control. TTC5 sg5 was used for all subsequent TTC5 KD studies. (**D**) Cell proliferation across early differentiation (iPSC to D3 i3N) quantified as number of Halo-NLS nuclei for NT (gray) and TTC5 KD (teal) cells; n = 10 wells/condition. Error bars, mean +/-SD; individual lines represent two independent experiments. (**E - F**) KD efficiency for TTC5, SCAPER and CCR4-Not1 complex components CNOT10 and CNOT11 in D14 i^3^Neurons at the (**E**) RNA level by RT-qPCR, normalized to HPRT and NT control, and at the (**F**) protein level by Western blot, with GAPDH control. (**G**) Raw transcript counts for all expressed (> 0 count) α-and β-tubulin isoforms obtained from RNA-Seq of untreated NT and TTC5 KD D14 i3Ns shown in Figure 1D.

**Figure S2.**
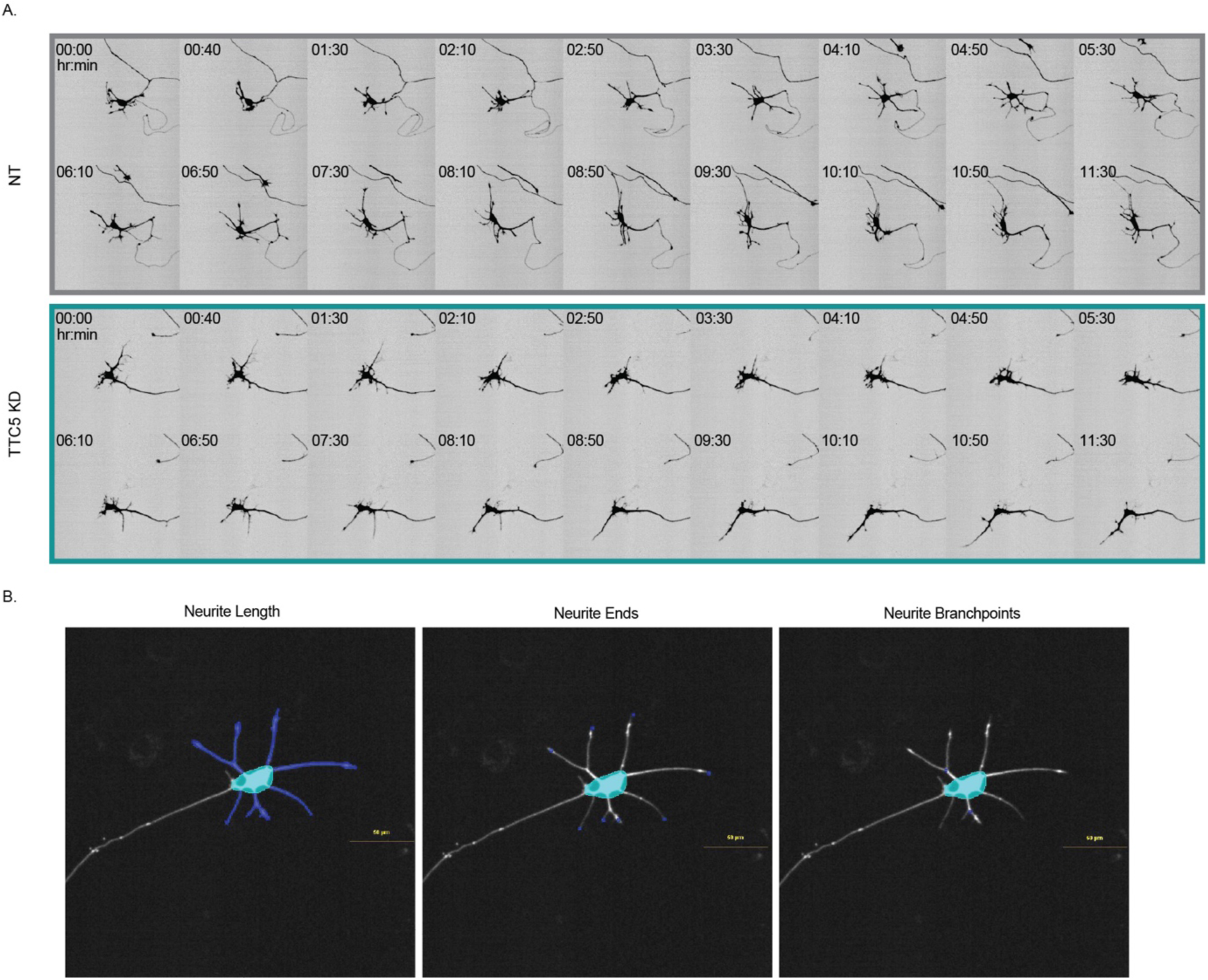
Representative examples of image collection and automated analysis of arborization metrics (related to Figure 2). (**A**) Montage of frames (every fourth frame) of mScarlet-expressing D12 i3Ns for arborization analysis presented in Figure 2A illustrating dynamic extension and retraction of processes for individual neurons (also see Movie S1). (**B**) Snapshots from customized Nikon GA3 automated pipeline (Methods) highlighting identification in dark blue, from left to right, of neurites (with omission of longest axonal process), neurite ends and neurite branchpoints.

**Figure S3.**
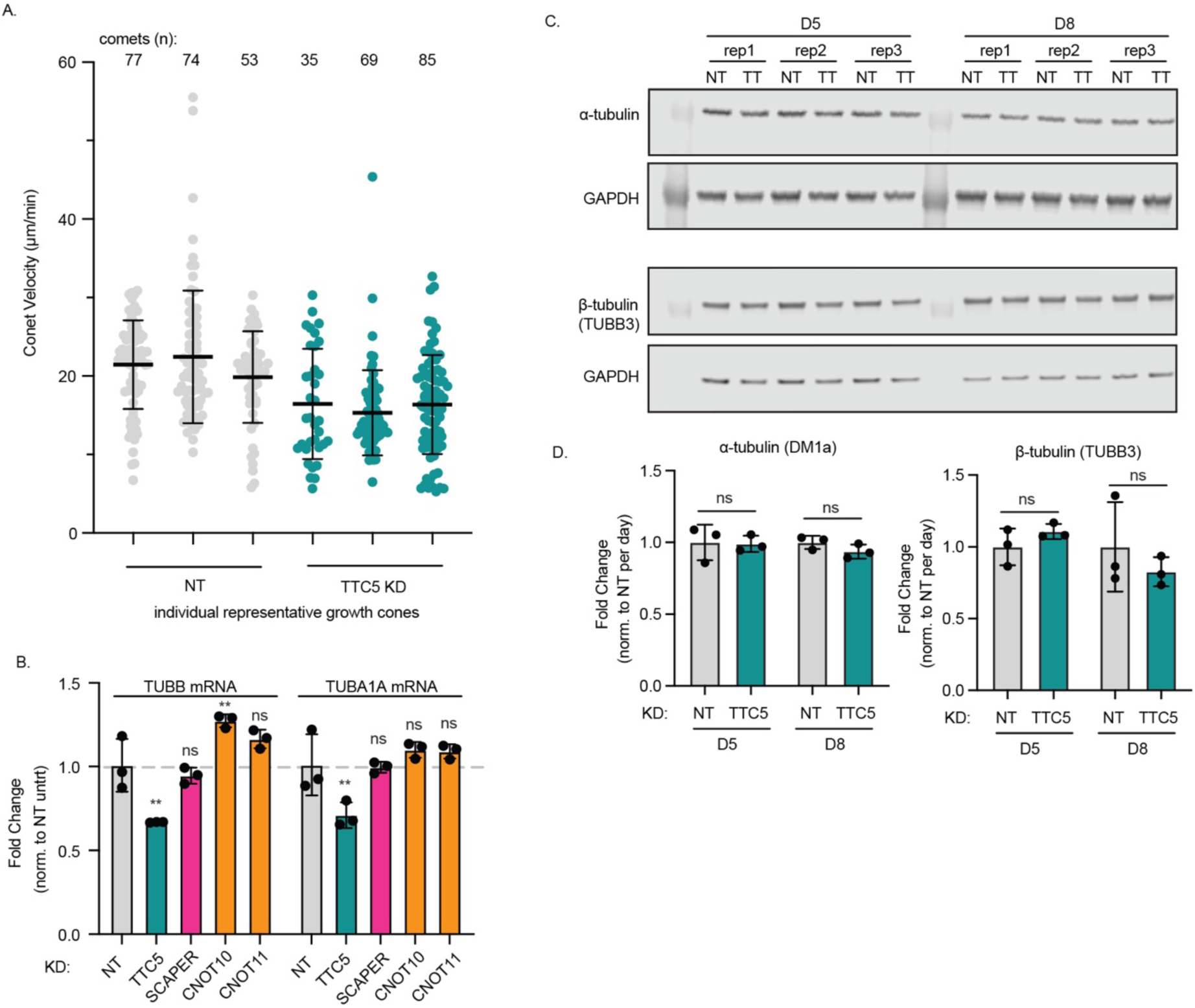
Additional characterization of EB1 comet dynamics and tubulin expression levels upon TTC5 KD (related to Figure 2). (**A**) EB1-GFP comet velocities for three representative individual growth cones for NT and TTC5 KD used to obtain average comet velocities per growth cone in Figure 2F. Error bars, mean +/-SD. (**B**) TUBB and TUBA1A mRNA levels assessed by RT-qPCR in D14 i3Ns upon KD of indicated gene normalized to NT control (following normalization to HPRT); error bars indicate mean +/- SD ; **, p < 0.01, by one-way ANOVA with Dunnett’s multiple comparisons test. (**C**) Western blot for α-tubulin and β-tubulin (TUBB3) tubulin protein in NT and TTC5 KD neurons at D5 and D8 in triplicate and quantified in (D) normalized to GAPDH; error bars indicate mean +/- SD; significance determined by unpaired t-test.

**Figure S4.**
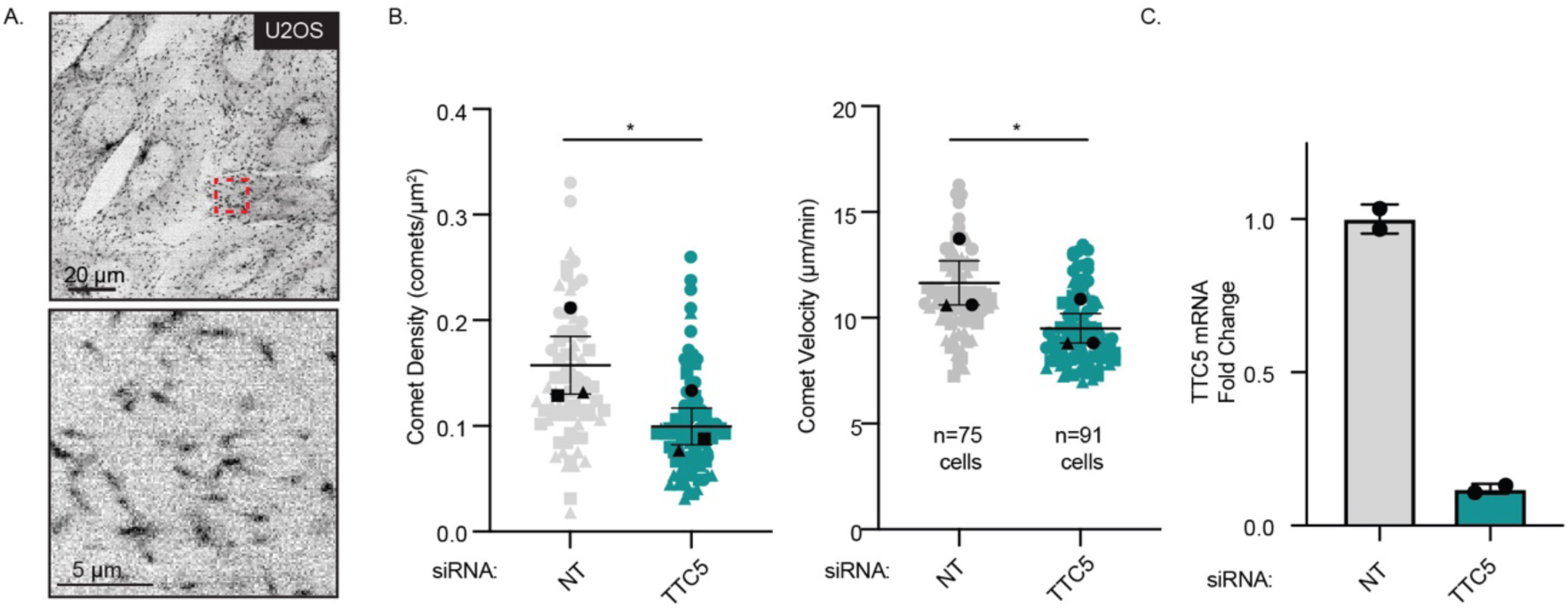
TTC5 loss results in reduced microtubule density and growth rate in U2OS cells (related to Figure 2). (**A**) Representative frame from a movie of EB1-mNG knock-in U2OS cells (top; scale bar: 20 μm) and magnified region of an U2OS cell showing individual EB1-mNG comets (bottom; scale bar: 5 μm). (**B**) EB1 comet density (left) and velocity (right) for NT (grey) or TTC5 siRNA KD (teal) cells. Individual points represent average values for individual cells from three independent experiments; n= 75, 91 cells for NT and TTC5 KD, respectively; error bars and symbols indicate the paired means of each independent experiment +/- SEM; *, p < 0.05 by paired t-test. (**C**) TTC5 mRNA level upon siRNA KD in U2OS cells assessed by RT-qPCR normalized to NT control (following normalization to GAPDH).

**Figure S5.**
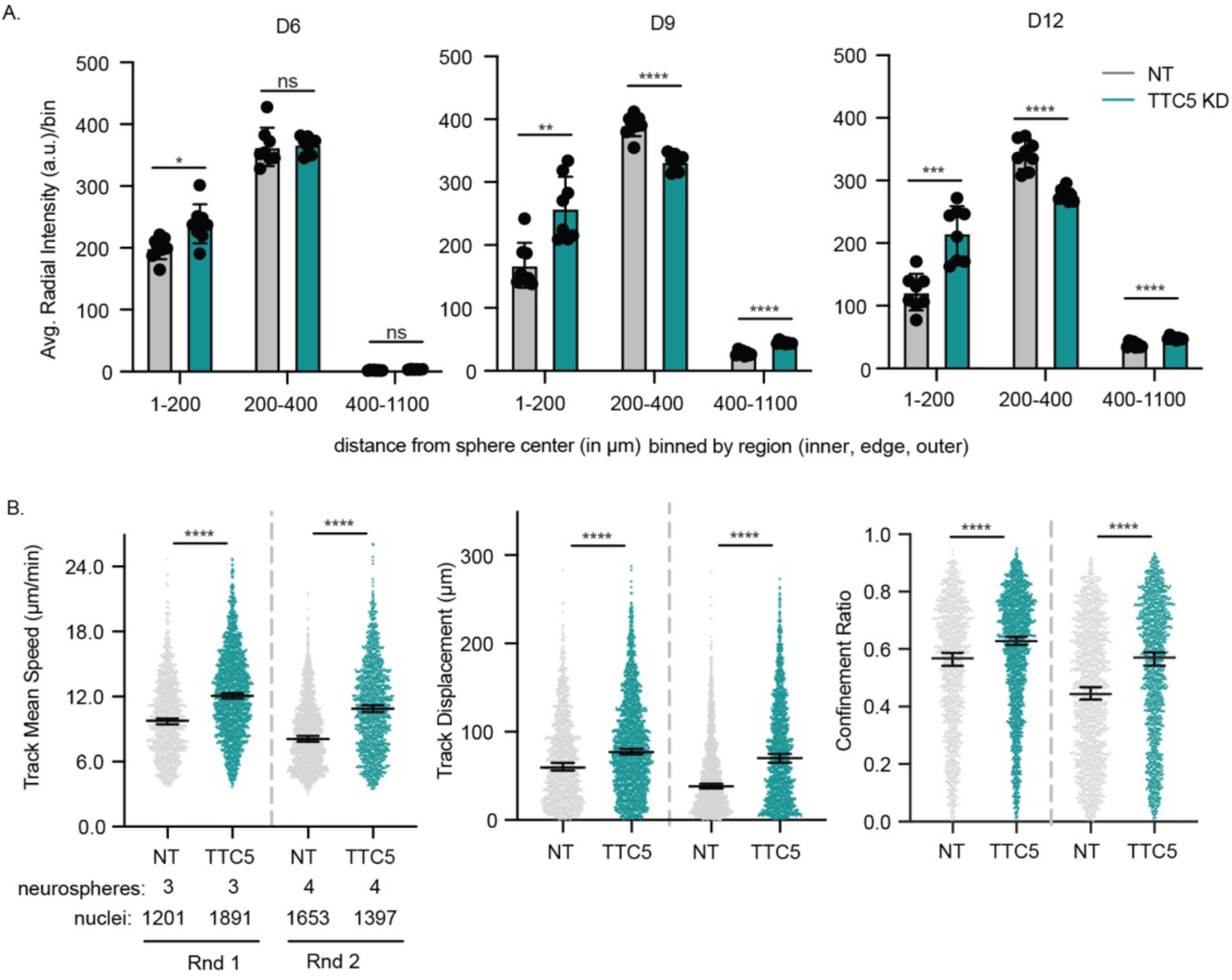
Alternate graphical depiction of static and dynamic motility readouts in Figure 3 (related to Figure 3). (**A**) For each timepoint (D6, D9, D12), the average signal of dispersed nuclei was divided into bins based on relative distance from the center and edge of each neurosphere (as in Figure 3B, right panel). Statistical significance was determined for the average signal from each bin; corresponding with the continuous plots depicting nuclear signal intensity at distances from the nucleus displayed in Figure 3B. Error bars indicate mean Halo-tagged nuclear signal within each indicated bin +/- SD, statistical significance determined by unpaired t-test (NT= 8 neurospheres, TTC5 KD=8 neurospheres); *, p<0.05, **, p < 0.01, ***, p < 0.001, ****, p < 0.0001. (**B**) Scatterplot of data represented as CDFs in Figure 3D of nuclear motility parameters for NT versus TTC5 KD neurospheres (Rnd 2 used for representative depictions in main figure). Error bars indicate the median with 95% confidence interval for parameters of individual nuclei pooled across multiple neurospheres for two independent experimental replicates; ****, p < 0.0001 by Kolmogorov-Smirnov test.

**Figure S6.**
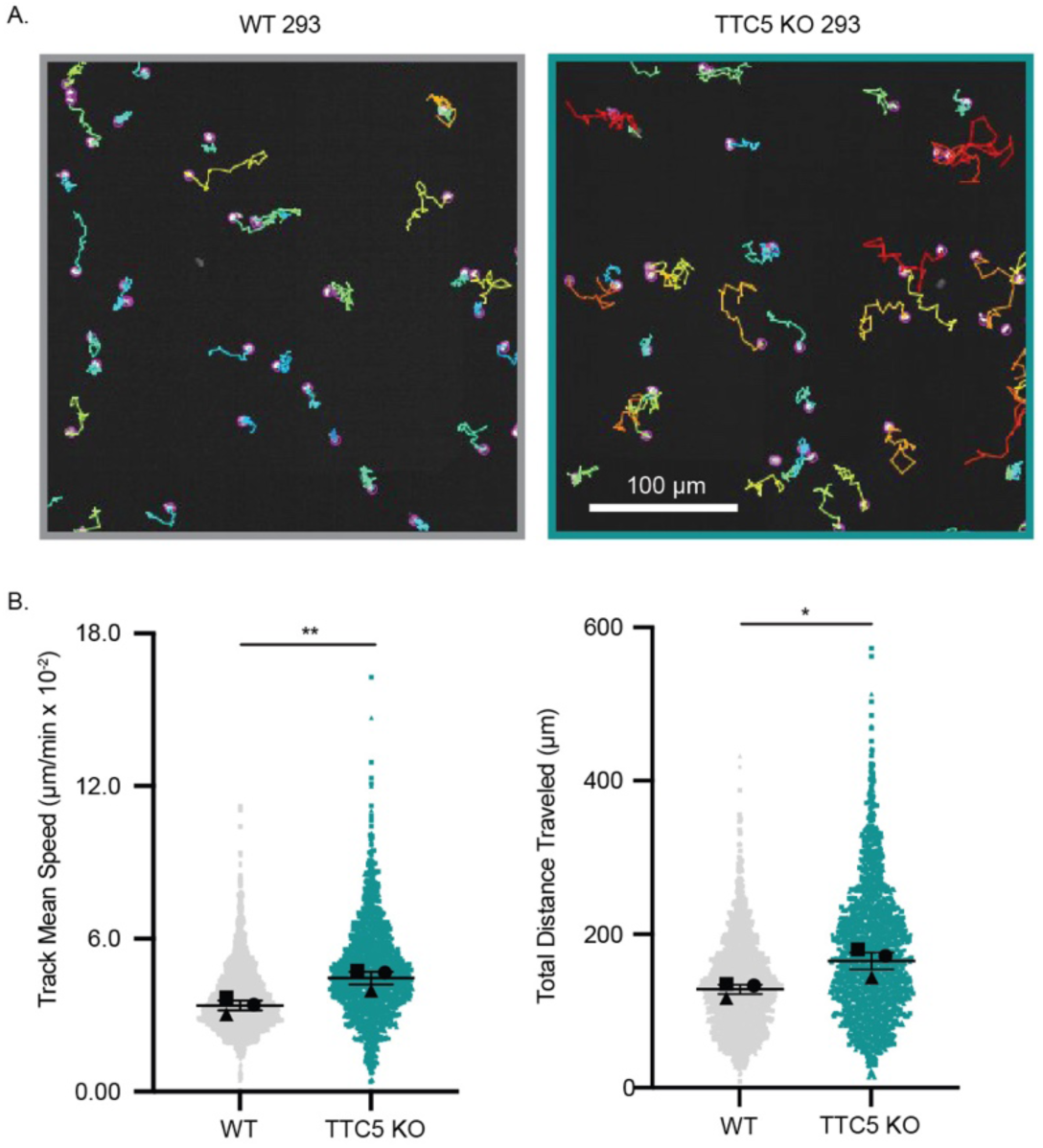
TTC5 loss results in altered motility in a non-neuronal cell line (related to Figure 3). (**A**)First frame from representative movie of Halo-tagged nuclei from WT and TTC5 KO HEK293 cells seeded onto fibronectin-coated plates (Methods). Nuclei are overlaid with tracks across a full eight-hour timecourse and color-coded by track mean speed from dark blue (0.00 mm/sec) to red (0.015 mm/sec). (**B**) mean speed (left) and total distance traveled (right) of tracked Halo-NLS nuclei. Dots represent individual nuclei compiled across three independent experiments (n= 1855 nuclei for WT, 1724 nuclei for TTC5 KO) with error bars and symbol indicating the paired medians of each of these three independent experiments +/- SEM; *, p < 0.05, **, p < 0.01 by paired t-test.

**Figure S7.**
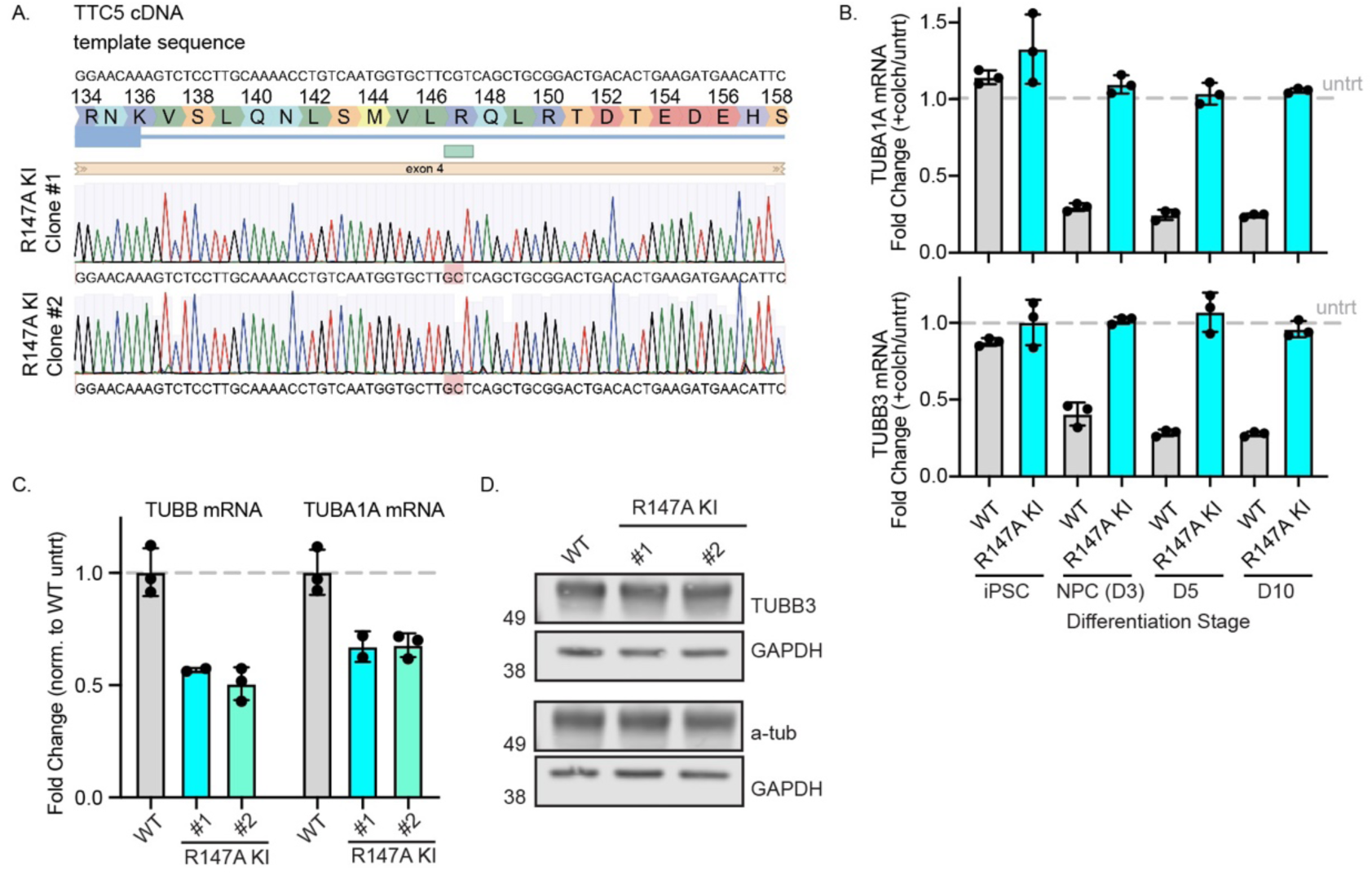
Line validation, tubulin autoregulation and tubulin level characterization in R147A KI lines (related to Figure 4). (**A**) Sanger sequencing validation of R147A KI clones #1 and #2. (**B**) Fold change for TUBB3 and TUBA1A mRNA level upon colchicine treatment relative to the untreated control for WT (gray) and R147A KI #1 (cyan) following initial normalization to housekeeping gene HPRT across differentiation from iPSCs to D10 i^3^Neurons assayed by RT-qPCR. Each condition was assayed in triplicate. (**C**) Fold change of tubulin mRNA levels relative to the WT control in untreated D14 WT and R147A KI i3Ns assayed by RT-qPCR as in (B). (**D**) Quantification of tubulin protein levels by Western blot in untreated D14 WT and R147A KI i^3^Neurons.

**Figure S8.**
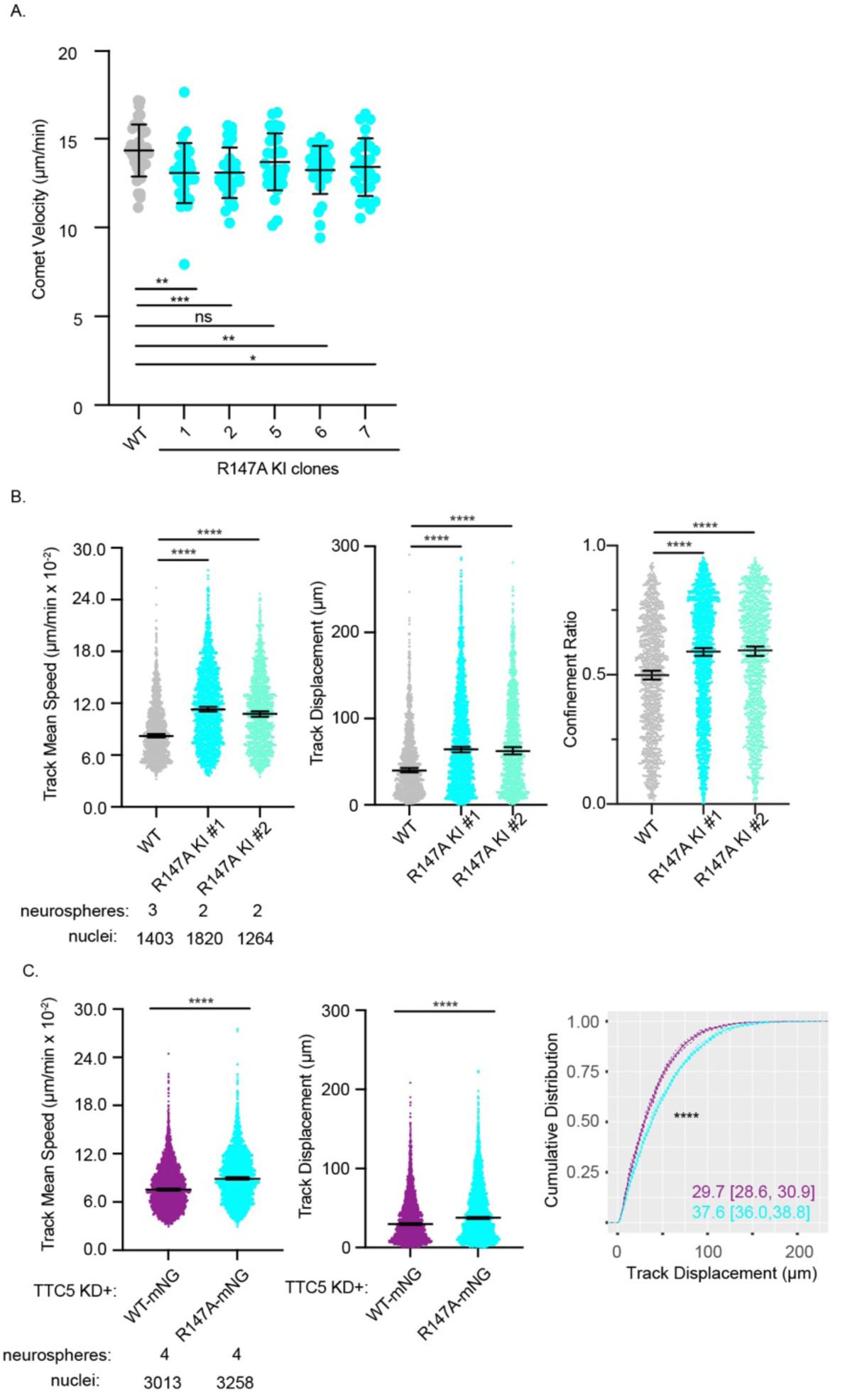
Additional characterization of neuronal phenotypes in the R147A knock-in and WT versus R147A mutant rescue i^3^Neurons (related to Figure 4). (**A**) Quantification of EB1-EGFP comet velocities across 5 independent R147A KI clones displaying mean +/-SD; *, p < 0.05, **, p < 0.01, ***, p < 0.001 by unpaired student’s t-test. (**B**) Scatterplot of data represented as a CDF in Figure 4H for nuclear motility parameters. Error bars indicate the median with 95% confidence interval for individual nuclei pooled across multiple neurospheres; ****, p < 0.0001 by Kruskal-Wallis test. (**C**) Scatterplot of data represented as CDF in Figure 4L (track mean speed) or in rightmost panel (track displacement), of nuclear motility parameters for TTC5 KD neurospheres upon reintroduction of either WT (purple) or R147A (cyan) TTC5-mNG. Error bars indicate the median with 95% confidence interval for parameters of individual nuclei pooled across multiple neurospheres; ****, p < 0.0001 by Kolmogorov-Smirnov test.

**Figure S9.**
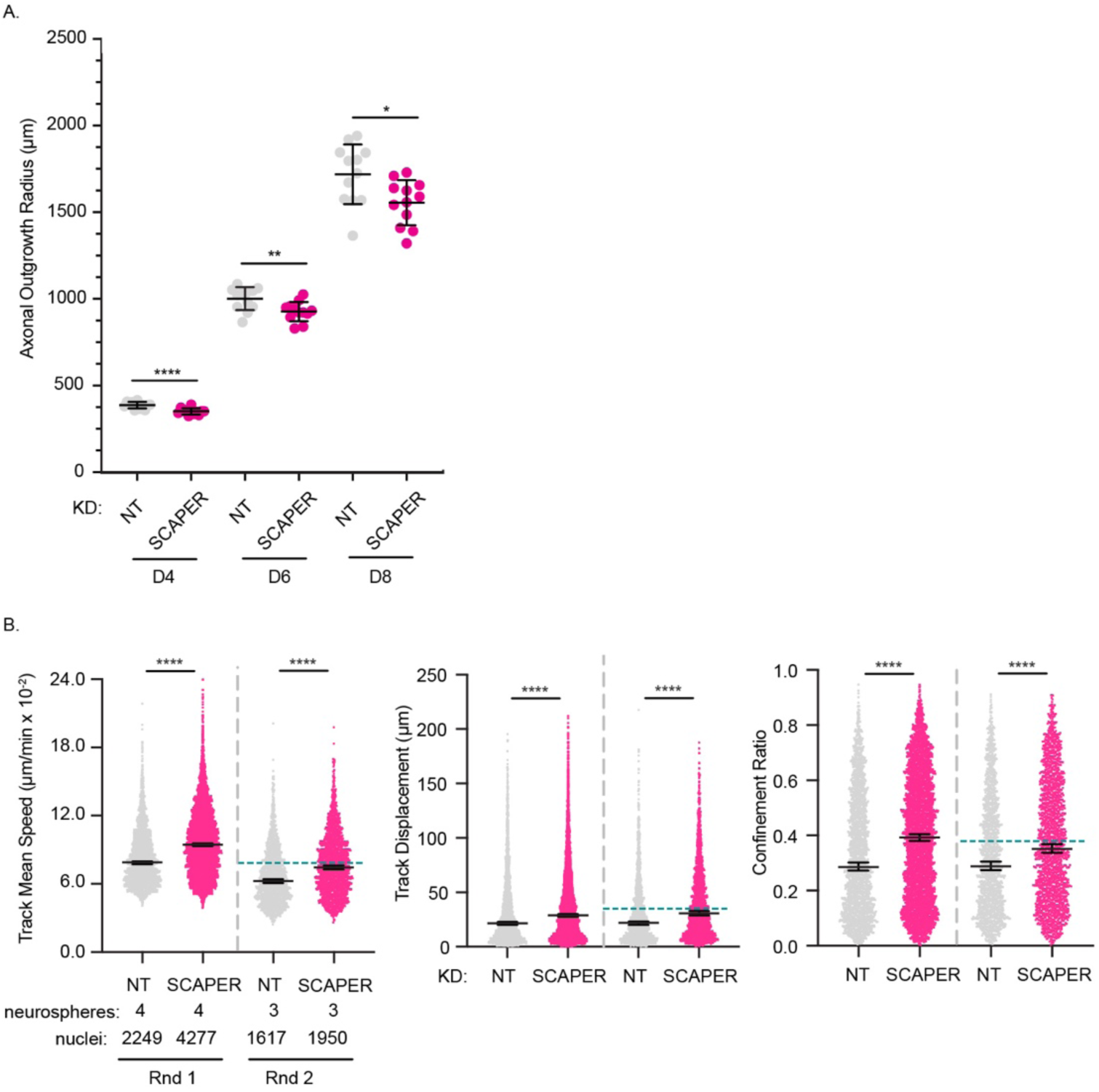
Additional neuronal assay characterization of SCAPER KD i^3^Neurons (related to Figure 5). (**A**) Independent experimental replicate of axonal outgrowth in NT versus SCAPER KD i3Ns assayed longitudinally at D4, D6 and D8 post-differentiation (n=12 spheres NT and 12 spheres SCAPER KD) as in Figure 5C. Error bars indicate the mean +/- SD with statistical significance determined by Welch’s t-test; *, p < 0.05, **, p < 0.01, ****, p < 0.0001 . (**B**) Scatterplot of data shown as CDF in Figure 5D, tracking nuclear motility parameters. Error bars indicate the median with 95% confidence interval for parameters of individual nuclei pooled across multiple neurospheres for two independent experimental replicates, ****, p<0.0001 by Kruskal-Wallis test. Dotted teal line in Rnd 2 replicate indicates respective median value for TTC5 KD neurospheres (n = 2) collected side-by-side within the same experiment.

**Figure S10.**
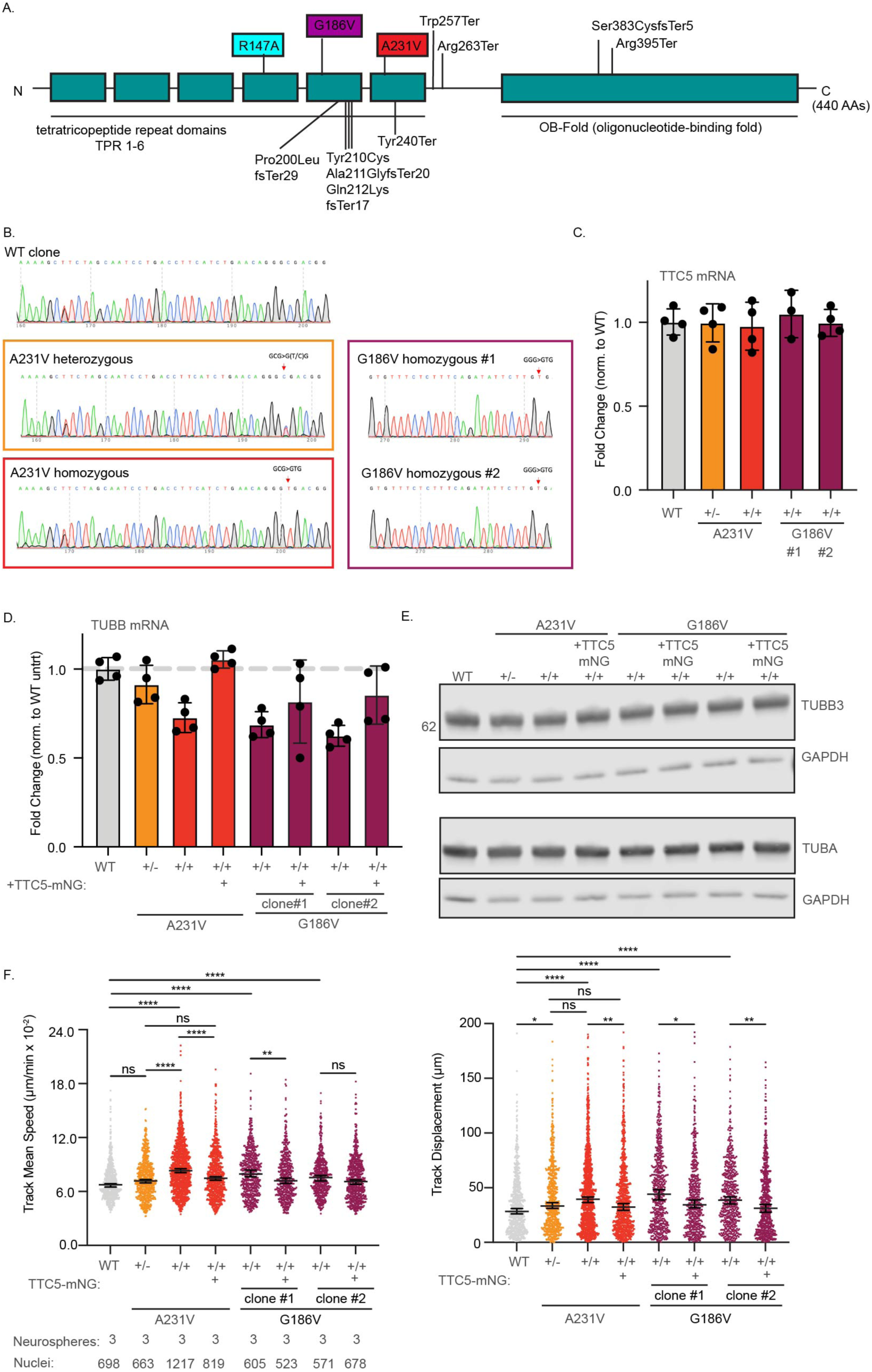
Baseline characterization and neuronal assay assessment for disease mutant KI lines (related to Figure 6). (**A**) Schematic of reported disease mutants for TTC5 with KI mutants analyzed in this study highlighted alongside structure-guided mutant R147A (modified from Musante et al.^18^). (**B**) Validation of disease mutant knock-ins via Sanger sequencing (**C**) Validation of disease mutant TTC5 mRNA levels by RT-qPCR (normalized to WT clone following normalization to HPRT) (**D**) Baseline TUBB mRNA level in untreated WT and disease mutant knock-in D10 i^3^Neurons by RT-qPCR (normalized to WT clone following normalization to HPRT). Each condition was performed in duplicate across two independent experiments. (**E**) Expression of α and β-tubulin protein in NT control and indicated disease mutant D10 i^3^Neurons assayed by Western blot (with GAPDH control). (**F**) Scatterplot depiction of data represented as CDF in Figure 6D (with inclusion of an additional G186V clone) of nuclear motility parameters track mean speed and track displacement. Error bars indicate the median with 95% confidence interval for individual nuclei pooled across neurospheres (n=3 per condition); *, p < 0.05, **, p < 0.01, ***, p < 0.001, ****, p < 0.0001 by Kruskal-Wallis test.

## Supplemental Movies

**Movie S1**. Time lapse showing dendritic extension dynamics and automated analysis for NT and TTC5 KD i^3^Neurons (related to Figure 2A, 2B).

**Movie S2**. Time lapse of EB1-GFP in NT and TTC5 KD i^3^Neurons growth cones (related to Figure 2E,2F).

**Movie S3.** Time lapse of migration for NT and TTC5 KD i^3^Neurons on glial bed (related to Figure 3C, 3D).

**Movie S4**. Time lapse of migration for WT and TTC5 R147A KI i^3^Neurons on glial bed (related to Figure 4H).

